# Declines in ice cover induce light limitation in freshwater diatoms

**DOI:** 10.1101/2023.08.02.551672

**Authors:** Brittany N. Zepernick, Emily E. Chase, Elizabeth R. Denison, Naomi E. Gilbert, Robbie M. Martin, Alexander R. Truchon, Thijs Frenken, William R. Cody, Justin D. Chaffin, George S. Bullerjahn, R. Michael L. McKay, Steven W. Wilhelm

**Author notes:** **Correspondence:** Steven W. Wilhelm, 1-865-974-0665, Robert. M. McKay, 1–.

## Abstract

The rediscovery of diatom blooms embedded within and beneath Lake Erie ice cover (2007-2012) ignited an intense interest in psychrophilic adaptations and winter limnology. Subsequent studies determined ice plays a vital role in winter diatom ecophysiology, as diatoms partition to the underside of ice thereby fixing their location within the photic zone. Yet, climate change has led to widespread ice decline across the Great Lakes, with Lake Erie presenting a nearly ice-free state in several recent winters. It has been hypothesized the resultant turbid, isothermal water column will induce light limitation amongst winter diatoms, serving as a detrimental competitive disadvantage. Here, we conducted a physiochemical and metatranscriptomic survey of the winter Lake Erie water column (2019-2020) that spanned spatial, temporal, and climatic gradients to investigate this hypothesis. We determined ice-free conditions decreased diatom bloom magnitude and altered diatom community composition. Diatoms increased the expression of various photosynthetic genes and iron transporters, suggesting they are attempting to increase their quantity of photosystems and light-harvesting components (a well-defined indicator of light limitation). Notably, we identified two gene families which serve to increase diatom fitness in the turbid ice-free water column: proton-pumping rhodopsins (a second means of light-driven energy acquisition) and fasciclins (a means to “raft” together to increase buoyancy and co-locate to the surface to optimize light acquisition). With large-scale climatic changes already underway, our observations provide insight into how diatoms respond to the dynamic ice conditions of today and shed light on how they will fare in a climatically altered tomorrow.

## Introduction

Winter was historically considered a period of planktonic persistence rather than growth [1, 2]. Yet, a series of limnological surveys conducted in the winters of 2007-2012 contested this with the rediscovery of dense diatom blooms associated with ice cover of Lake Erie (US, Canada) [3, 4]. This finding ignited interest in winter limnology [5–7], with subsequent studies demonstrating ice-associated communities were dominated by the centric colonial diatoms *Aulacoseira islandica* and *Stephanodiscus* spp. [3, 4, 8–10]. Notably, chlorophyll *a* (Chl *a*) concentrations during winter surpassed those of spring [3] and examinations of silica deposition in frustules demonstrated cells were metabolically active [11]. Additional studies emphasized these blooms are of biotic and biogeochemical importance, as winter-spring diatom biovolumes surpass summer cyanobacterial biovolumes by 1.5- to 6-fold [12] and drive recurrent summer hypoxia in the Lake Erie central basin [7, 10, 12]. Though Lake Erie serves as a leading case study for winter diatom blooms, they are far from an isolated phenomenon. Blooms are often unreported due to a lack of winter surveys [13], yet they have been well-documented beneath the ice in Lake Baikal [14, 15] and characterized in other north temperate freshwater systems such as The Loch (US) [16], Lake Barleber (Germany) [13], Lakes Ladoga and Onega (Russia) [17], Lake Kasumigaura (Japan) [18] and the River Danube (Hungary) [19].

Contributing to the ecological success of winter diatoms are adaptations that increase membrane fluidity and enhance light harvesting in icy, low-light conditions [9]. Yet, arguably a major adaptation responsible for winter diatom success is their ability to partition to surface ice cover *via* interactions with ice-nucleating bacteria, which allows diatoms to co-locate themselves to the under-ice surface to maintain an optimal light climate for photosynthesis [8, 20]. Cumulatively, studies demonstrate ice cover plays a critical role in shaping winter diatom ecophysiology and increasing competitive fitness in the icy water column [3, 4, 8, 9, 11, 20]. Yet, this raises the question of how this keystone phyla [21–24] will fair in a climatically altered ice-free future.

Lake Erie, concomitantly with other lakes across the globe, is experiencing unprecedented declines in ice cover [7, 25, 26]. Notably, projections suggest ice cover may disappear entirely across the Great Lakes by the end of the century [27]. This loss of ice cover presents a unique scenario for shallow lakes such as Erie (mean depth ∼19 m). Due to predominant westerly winds blowing across the lakes west-to-east axis, snow seldomly accumulates on the surface ice [3, 20, 28] (Figure 1A), allowing light to penetrate below ice cover where diatoms are located. Yet, in the absence of ice cover, winds create an isothermal turbid water column in this shallow lake [29, 30] (Figure 1B). Indeed, Beall et al., [4] reported turbidities (NTU) a magnitude higher in ice-free Lake Erie (2012) compared to the ice-covered year prior (2011), and noted diatom abundances significantly declined in the turbid water column. This study suggested light limitation was the key driver of diatom decline, citing elevated turbidity within an ice-free lake would induce light limitation [4, 7, 31, 32].

**Figure 1:**
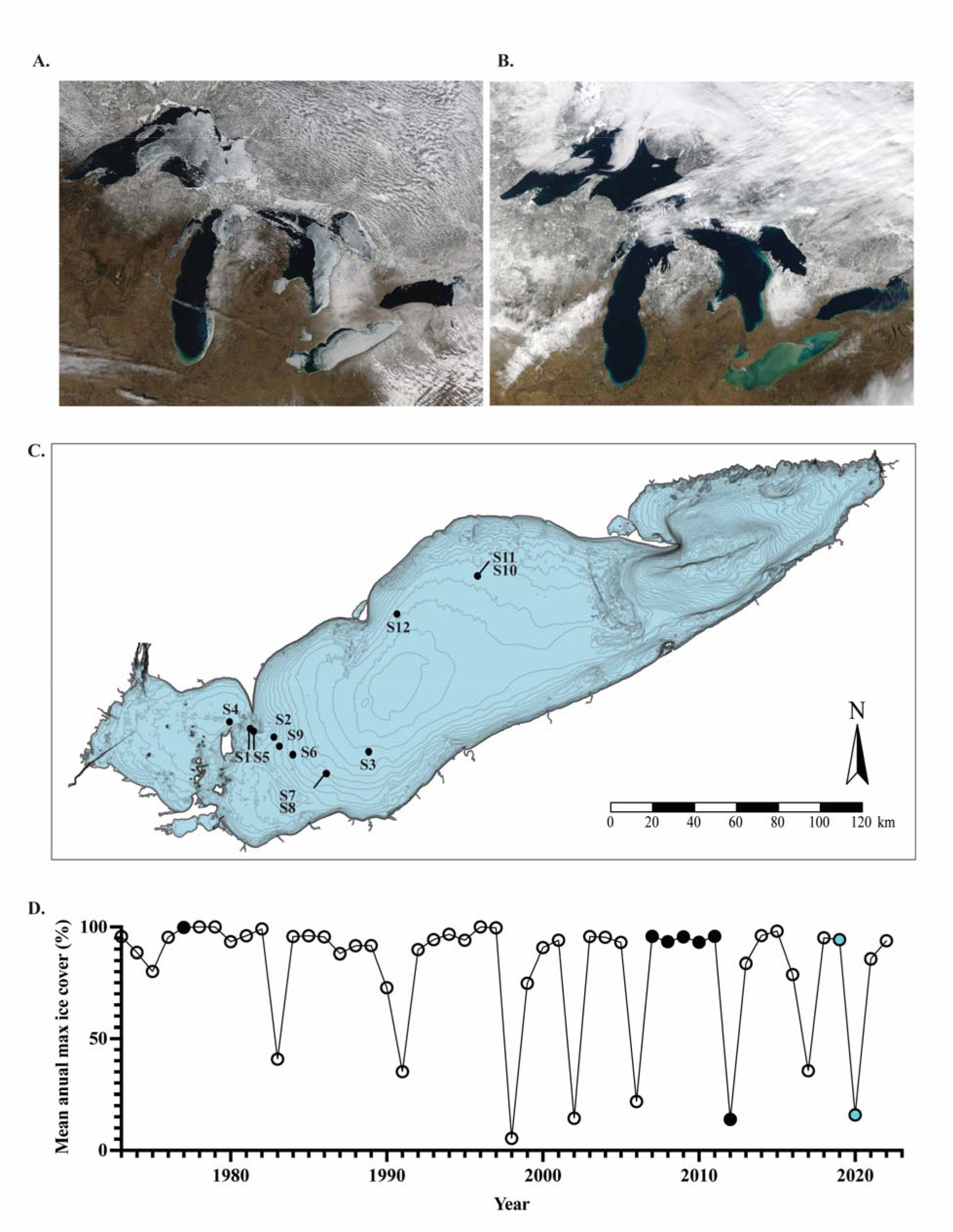
Climatic, spatial and temporal variability across Lake Erie samples. **(A)** MODIS satellite image (March 16^th^, 2014) depicting a large amount of ice cover across the Great Lakes. During the winter of 2014, Lake Erie had a mean annual ice cover of ∼80%. **(B)** MODIS satellite image (February 12^th^, 2023) depicting a lack of ice cover across the Great Lakes. Sediment plumes can be observed throughout Lake Erie. During the winter of 2023, Lake Erie had a mean annual ice cover of ∼8%. Photo Credit: NOAA GLERL/NOAA Great Lakes CoastWatchNode. **(C)** Sample sites across Lake Erie visited throughout winter-spring 2019 and 2020. **(D)** Historical trends in Lake Erie mean annual maximum ice cover (%). Open circles are years that (to our knowledge) do not have peer-reviewed published planktonic survey data. Solid black circles are years that were previously surveyed in prior published studies. Solid blue circles are years sampled in this study. Figure adapted from data retrieved from NOAA GLERL database [96].

We employed *in situ* analyses and metatranscriptomics to explore the hypothesis that winter diatoms are light limited in the ice-free water column. Facilitated by collaborative efforts with the U.S. and Canadian Coast Guards [33], opportunistic samples were collected throughout 2019 and 2020, yielding winter samples collected from both ice-covered (2019) and ice-free (2020) water columns [34]. The survey also included spring samples (outgroup). Our analyses confirm ice cover alters diatom bloom magnitude and phylogeny while providing novel evidence of light limitation within diatom communities of the ice-free water column. We also present evidence of two adaptations which we hypothesize increase the competitive fitness of freshwater diatoms within the ice-free winter water column.

## Methods

### Lake Erie winter-spring water column sampling

Samples of opportunity (n = 77) from the Lake Erie planktonic community were collected across temporal, spatial, and climatic gradients throughout the winter of 2019 and 2020. This large-scale collaborative effort included multiple surveys conducted by USCGC *Neah Bay*, CCGS *Limnos* and M/V *Orange Apex,* resulting in a large metatranscriptomic dataset [34]. Prior to sample collection, water column physiochemical parameters were recorded along with meteorological conditions and ice cover observations. Briefly, water samples were collected from 0.5 m below the surface and processed for analyses of dissolved and particulate nutrients (mg L^-1^), size-fractionated (< 0.22-µm and < 20 µm) Chl *a* biomass (µg L^-1^), phytoplankton taxonomy and enumeration (cells L^-1^), and total community RNA. Metadata are available online at the Biological and Chemical Oceanography Data Management Office (BCO-DMO) [35]. Refer to Supplemental Methods/Results and Supplemental Tables for further details.

### RNA extraction and sequencing

RNA extractions were performed using previously described phenol-chloroform methods with ethanol precipitation [36]. Residual DNA in samples was digested *via* a modified version of the Turbo DNase protocol using the Turbo DNA-free kit (Ambion, Austin, TX, USA). Samples were determined to be DNA-free *via* the absence of a band in the agarose gel after PCR amplification using 16S rRNA primers as previously reported [34]. Samples were quantified using the Qubit RNA HS Assay Kit (Invitrogen, Waltham, MA, USA) and sent to the Department of Energy Joint Genome Institute (DOE JGI) for ribosomal RNA reduction and sequencing using an Illumina NovaSeq S4 2 × 151-nucleotide indexed run protocol (15 million 150-bp paired-end reads per library) as reported previously [34].

### Metatranscriptomic analysis

Filtering and trimming of raw reads was performed by DOE JGI using BBDuk (v.38.92) and BBMap (v.38.86) [37, 38]. Bioinformatic processing was conducted using a prior-established metatranscriptomic workflow [39]. Trimmed and filtered libraries (n = 77) were concatenated and assembled (co-assembled) using MEGAHIT (v.1.2.9) [40]. Co-assembly statistics were determined *via* QUAST QC (v.5.0.2) [41]. Trimmed reads were mapped to the co-assembly using BBMap (default settings) (v.38.90) [38]. Gene predictions within the co-assembly were called using MetaGeneMark (v.3.38) [42] using the metagenome style model. Taxonomic annotations of predicted genes were determined using the MetaGeneMark protein file, EUKulele (v.1.0.6) [43] and the PhyloDB (v.1.076) database. We note, this study brings to light a challenge within the freshwater field at large: there is a lack of sequenced freshwater diatom taxa and an absence of freshwater taxonomic annotation databases, which constrains the taxonomic resolution of bioinformatic data. Indeed, in 2016, Edgar et al., [9] noted only 23% of the taxonomic diatom annotations within their Lake Erie metatranscriptome could be tied to genera known to be present within the Great Lakes. Earlier this year, Reavie [44] reiterated the lack of studies regarding Great Lakes diatoms, citing there are various undescribed and unclassified diatom taxa to date. As a result, there may be transcriptional changes within the winter diatom community which have gone undetected within this study, specifically at the genera level. Many of the taxonomic annotations we generated were best aligned to marine counterparts due to a lack of sequenced freshwater representatives (*i.e.,* where reads are annotated as belonging to a genus “*-like*” genome). More broadly, sequence data is often not coincident with classic morphological taxonomy. Nonetheless, *A. islandica* filaments exhibit a distinct morphology from *Stephanodiscus* spp., and bioinformatic pipelines such as EUKulele accurately distinguish the taxonomy of these “unsung eukaryotes” at the class level [43]. Thus, this ambiguity in diatom taxonomy does not negate the definitive disparity in diatom cell counts and transcription observed in this study. Following, genes were functionally annotated using eggNOG-mapper using a specified e-value of 1e^-10^ (v.2.1.7) [45]. Ensuing, featureCounts [46] within the subread (v.2.0.1) package was used to tabulate read counts to predicted genes. Mapped reads were normalized to TPM (Transcripts Per Million), representing relative “expression” values prior to statistical analyses (ANOSIM, SIMPER, nMDS etc.). To investigate transcriptional patterns of the winter diatom bloom community, we focused on a subset of libraries (n = 20) selected for consistency in sample collection methods (whole water filtration) and diatom abundances (Supplemental Table 1). Thus, all data reported hereafter pertains to these 20 libraries. Raw data for all 77 transcriptomic libraries are available at the JGI Data Portal (https://data.jgi.doe.gov) under Proposal ID 503851 [34]. Refer to Supplemental Methods/Results and Supplemental Tables for further details.

### Phylogenetic analysis

A phylogenetic tree of fasciclin containing domains (proteins of interest) was produced using differentially expressed (DE) putative proteins identified from this study (n=18), domains recovered from the eggNOG orthology database and publicly available domains from NCBI [47]. A custom database was curated using all NCBI diatom proteins. A DIAMOND (v.2.0.15) [48] blastp alignment was performed with putative fasciclin proteins and eggNOG domains against the diatom database to recover all putative diatom fasciclin domains. The recovered domains were then aligned (DIAMOND blastp) against the NCBI non-redundant database. These results were compiled and collapsed to 80% similarity using CD-HIT (v.4.7) [49] and a multiple sequence alignment was performed using MAFFT (v.7.310) [50] with 500 iterations. Gaps were closed using trimAl with gappyout (v.1.4.rev15) [51] and examined using AliView (v.1.28) [52]. A phylogenetic tree (1000 bootstraps) was constructed using a model test selecting for a general non-reversible Q matrix model estimated from Pfam database (v. 31) [53] with a gamma rate heterogeneity. The consensus tree was visualized using iTOL [54].

A phylogenetic tree of diatom PPR containing domains (proteins of interest) was produced using DE (n = 2) and non-DE (n = 9) putative proteins identified from this study (total n=11), NCBI non-redundant database (nr) putative proteins were searched using baited study sequences identified as rhodopsin/rhodopsin-like/rhodopsin and with a diatom taxonomic designation by EUKulele via diamond blastp (v. 2.0.15), the NCBI nr database was queried again with these results. NCBI nr was also queried in the same way with potential freshwater diatom whole genomes with no suitable results. Retrieved amino acid sequences were collapsed at 100% using CD-HIT and aligned by MAFFT (v.7.520) with 1000 iterations. The alignment was then trimmed using trimAL (v.1.4. rev15) and the automated1 parameter. IQ-TREE (v. 2.2.0.3) was used to produce a consensus tree with 1000 bootstrap iterations using model test results (Q.pgam+G4 model). The resulting tree was modified from iTOL (v.6) visualization. Refer to Supplemental Methods/Results and Supplemental Tables for further details.

## Statistical analyses

Comparisons of water column physiochemical features by ice cover were made in Prism (v. 9.3.1) *via* two-tailed unpaired *t*-tests. Variability in expression (TPM) between transcriptomic libraries was assessed *via* ANalysis Of Similarities (ANOSIM) and Similarity Percentage (SIMPER) analyses using PRIMER (v.7) [55]. Bray-Curtis similarity and non-metric multi-dimensional scaling (nMDS) were performed in R. Differential expression (DE) of transcript abundance was performed using DESeq2 (v.1.28.1) [56]. Genes with an absolute log_2_ fold change (Log_2_FC) >2 and adjusted p-value of < 0.05 were considered differentially expressed. Z-scores reported in heat maps were calculated by heatmapper.ca (Clustering method: Average linkage, Distance measurement method: Pearson) [57]) using the DESeq2 variance stabilizing transformed values (VST) [57]. Refer to Supplemental Methods/Results for further details.

## Results

### Physiochemical profiles and winter community characterization

Samples were collected across 12 sites throughout the central basin of Lake Erie with true biological replication at a subset of stations (Figure 1C) (Supplemental Table 1). Temporally, the samples span February-March 2019 and February-June 2020, yielding 14 winter and 6 spring libraries. Climatically, the winter of 2019 was a year of high ice cover (mean maximum ice cover of 80.9%), whereas winter 2020 was a year of negligible ice cover (mean maximum ice cover of 19.5%) [58] (Figure 1D). Libraries 1-4 were collected during ice cover (ranging from 3-15 cm in thickness) while winter libraries 5-14 were collected during ice-free conditions. Winter lake surface temperatures ranged from ∼0-6 °C across sample sites (Supplemental Figure 1A). Overall, nutrient concentrations at ice-covered sites were not significantly different from ice-free sites save for nitrate (Supplemental Figure 1B-H). While not significant (p ≥ 0.13), the highest total Chl *a* concentrations (> 0.22 µm) coincided with ice cover (Figure 2A, B). The larger sized-fraction of phytoplankton contributed an average of 70% (+/- 27%) to total Chl *a* during ice cover and 50% (+/- 13%) in ice-free winter sites (Supplemental Figure 2), but the differences were not significant (p = 0.22). Cell concentrations of diatoms (Bacillariophyta) dominated the winter water column regardless of ice conditions, with other eukaryotic phytoplankton (*e.g.,* Chlorophyta, Cryptophyta, and Dinophyta) present at concentrations 1-2 orders of magnitude lower (Supplemental Figure 3). While Bacillariophyta concentrations decreased slightly at ice-free sites (p = 0.33), Dinophyta concentrations significantly increased (p = 0.03), with Cryptophyta and Chlorophyta exhibiting similar trends (p ≥ 0.05). Overall, centric diatoms (Mediophyceae, Coscinodiscophyceae) dominated the winter diatom community while pennate diatoms (Bacillariophyceae, Fragilariophyceae) were found at concentrations an order of magnitude lower (Figure 2C, D). Despite this dominance, centric diatoms demonstrated a decreasing trend in ice-free samples while pennate diatoms exhibited significant increases in ice-free samples (p = 0.03) albeit remaining at low abundances. Cell concentrations of the centric bloom formers *Stephanodiscus* spp. and *A. islandica* were highest during ice cover (Figure 2E, F). Notably, *Stephanodiscus* spp. concentrations were significantly higher than *A. islandica* in ice-covered samples (p = 0.03), yet not significantly greater than *A. islandica* in ice-free samples (p = 0.19) (Supplemental Figure 4). Further, while small centric diatoms (5-20 μm size) were not detected in ice covered samples, they were found to range from ∼300-3,000 cells L^-1^ in ice-free samples (Supplemental Figure 5). These small centric diatom taxa accounted for ∼83% of the winter diatom community at site 8, although they otherwise contributed an average of 26% to the total diatom community in ice-free samples (Supplemental Figure 6).

**Figure 2:**
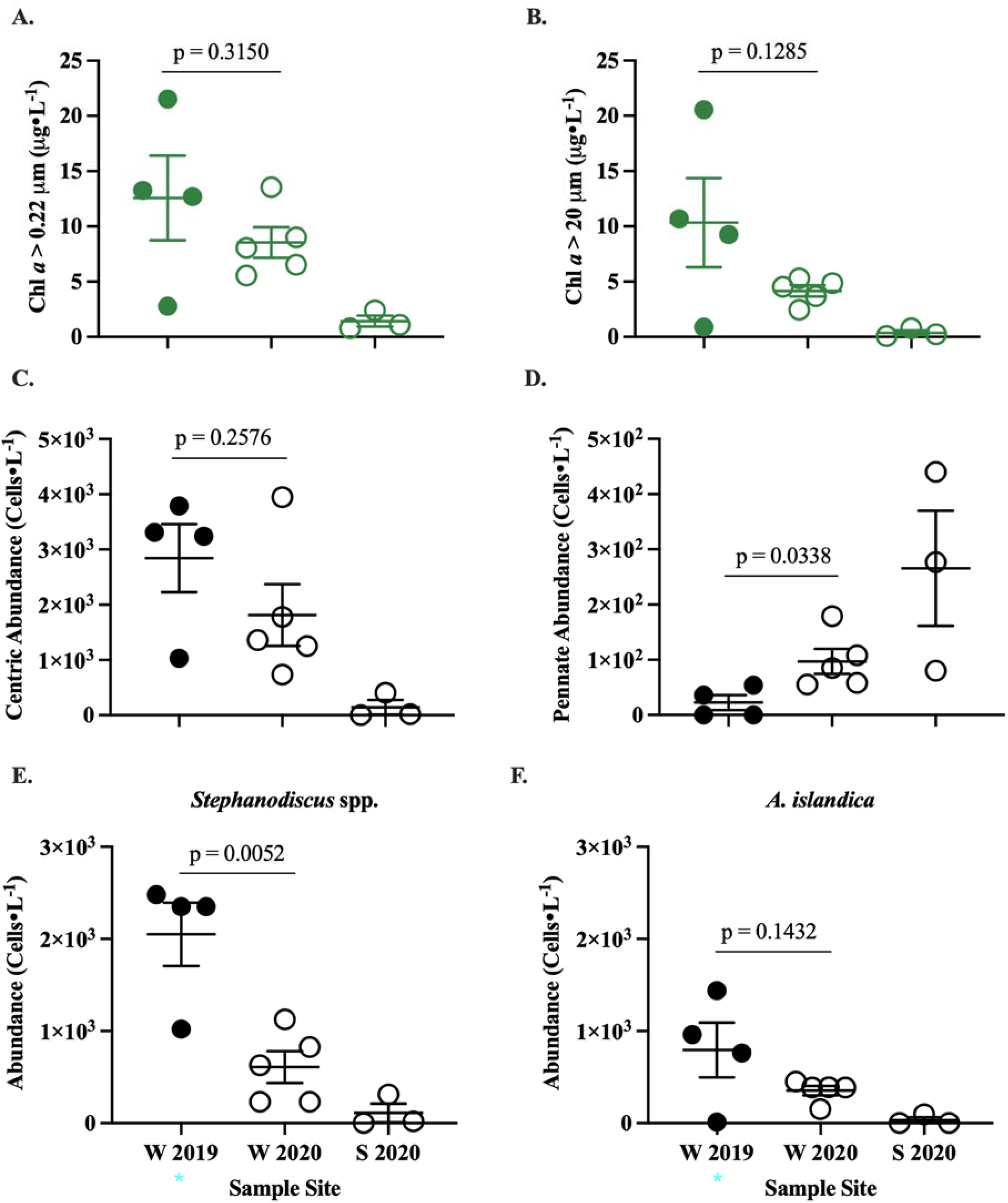
Characterization of biotic community across Lake Erie sample sites. Samples are organized on the x-axis by season (W = winter, S = spring) and year. Solid shapes indicate the sample was collected during ice cover (2019) open shapes indicate the sample was collected during no ice cover (2020). Ice cover samples are indicated by a blue asterisk. **(A)** Total Chl *a* concentration of the whole water column community (i.e., >0.22 μm in size) (μg L^-1^) **(B)** Chl *a* concentration of the large size fractioned community (i.e., >20 μm in size) (μg L^-1^). **(C)** Cell abundances (Cells ·L^-1^) of centric diatoms (*Stephanodiscus* spp. + *A. islandica* + Small centric diatoms of 5-20 μm). **(D)** Cell abundances of pennate diatoms (*Fragilaria* spp. + *Asterionella formosa* + *Nitzschia* spp). **(E)** Cell abundances (Cells ·L^-1^) of *Stephanodiscus* spp.,Mediophyceae class. **(F)** Cell abundances (Cells ·L^-1^) of *A. islandica,* Coscinodiscophyceae class.

### Transcriptomic response of winter diatom community to ice cover

Diatoms dominated the winter transcriptional pool across major eukaryotic phytoplankton communities regardless of ice cover (Figure 3A). In turn, diatoms of the class Mediophyceae dominated diatom community transcription regardless of ice cover (Figure 3B). At the genus level of each class, there was a lack of definitive trends across libraries thus they are omitted from the main text (Supplemental Figures 7-10). Overall, there was no correlation between diatom cell abundance and transcript abundance (Supplemental Figure 11). Refer to Supplemental Methods/Results and Supplemental Tables for further detail.

**Figure 3:**
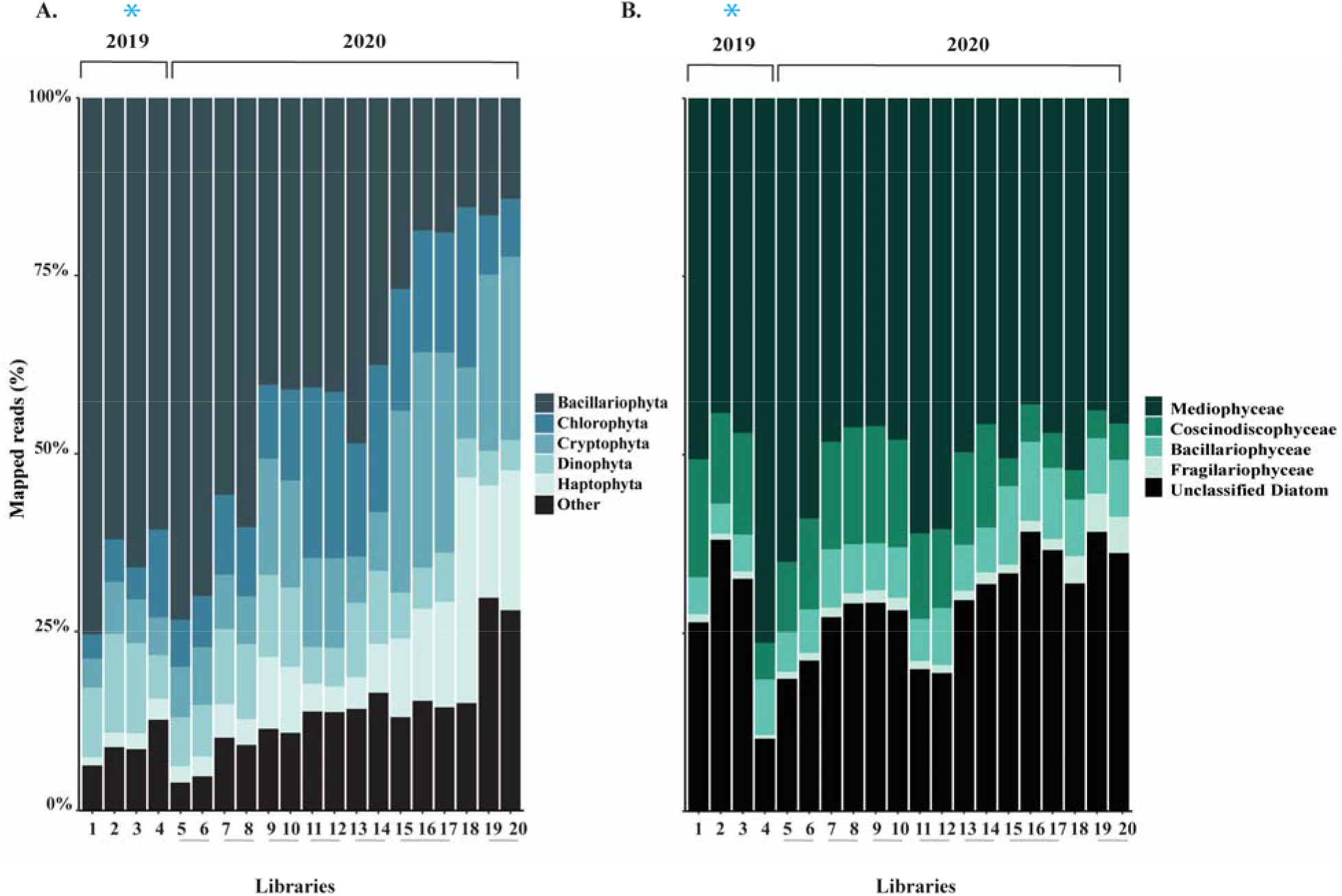
Relative transcript abundance of major eukaryotic phytoplankton taxa and diatom classes. Libraries are listed in chronological order of sample date on x-axes, with biological replicates joined by grey horizontal bars. Ice cover samples are indicated by a blue asterisk. **(A)** Relative transcript abundance of MEPT. All groups which formed <5% of the total mapped reads are included within “Other” (Amoebozoa, Hilomonadea, Excavata, Rhizaria, NA). **(B)** Relative transcript abundance of Bacillariophyta classes Mediophyceae, Coscinodiscophyceae, Bacillariophyceae, Fragilariophyceae and unclassified diatoms.

Normalized expression (TPM) profiles of the total water column community displayed clustering by ice cover (Figure 4A), with SIMPER analyses demonstrating an average dissimilarity of 64% between ice cover and ice-free winter libraries (Supplemental Table 1P). ANOSIM tests confirmed ice strongly affected winter community gene expression (R = 0.87, p = 0.002) (Supplemental Figure 12A) (Supplemental Table 1Q). Surprisingly, diatom community expression did not cluster as strongly by ice cover (Figure 4B), with SIMPER analyses indicating an average dissimilarity of 47% between ice cover and ice-free libraries (Supplemental Table 1T). ANOSIM tests confirmed ice cover exerts a lesser influence on winter diatom community expression overall compared to the full water column community (R = 0.282, p = 0.059) (Supplemental Figure 12B). In contrast, season had a strong effect on diatom expression (SIMPER Average dissimilarity = 77%; ANOSIM R = 0.927, p = 0.001) (Supplemental Tables 1V, W).

**Figure 4:**
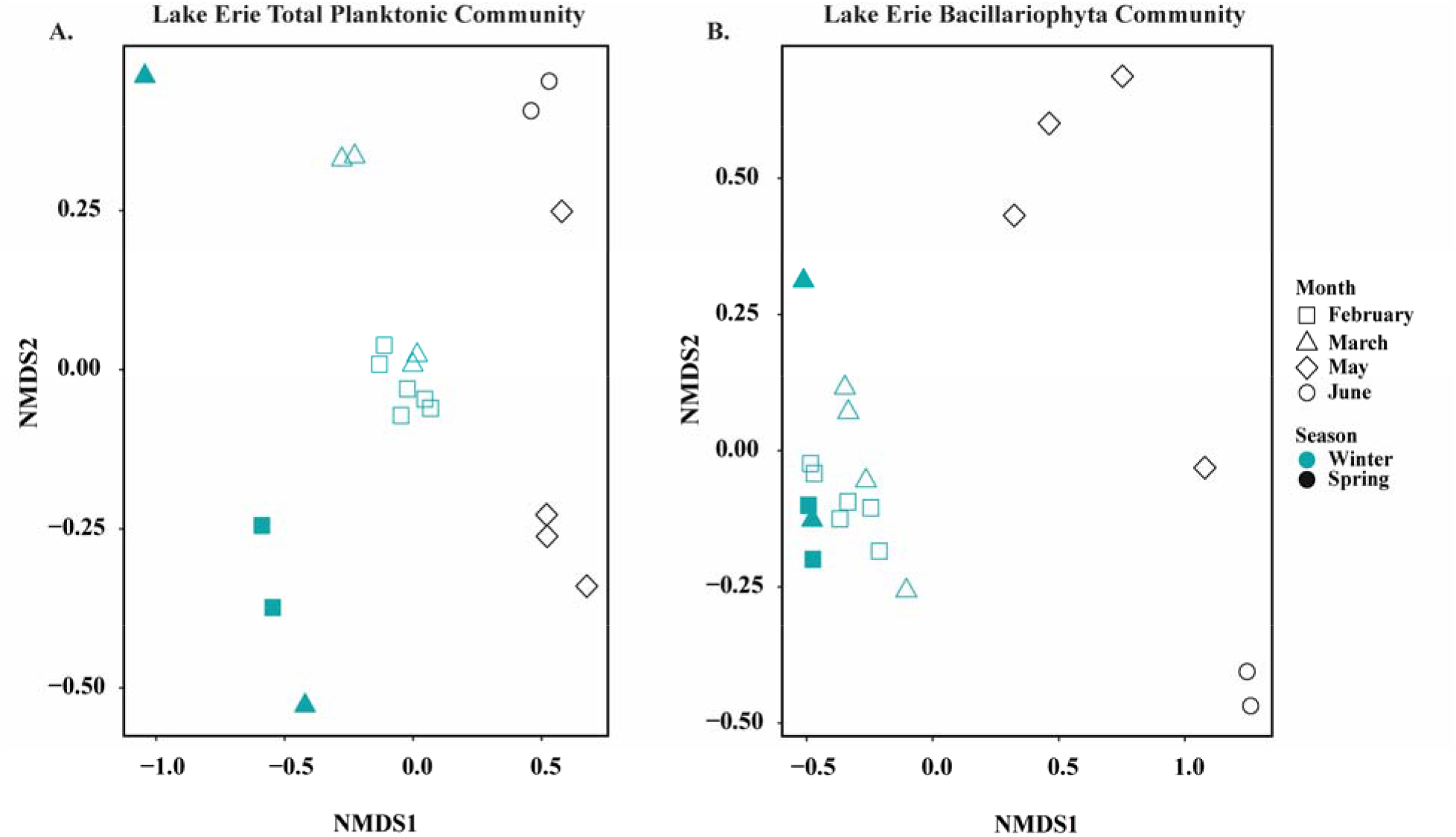
Similarity (Bray-Curtis) clustering of the 20 metatranscriptomic librariy normalized expression values (TPM). **(A)** nMDS of the entire water column community expression, stress value = 0.0633. (**B)** nMDS of the Bacillariophyta community expression, stress value= 0.0497. Samples are presented as follows: February = squares, March = triangles. May = diamonds, June = circles. Blue indicates the sample was collected during the winter, black indicates the sample was collected during the spring. Solid shapes indicate the sample was collected during ice cover (2019) open shapes indicate the sample was collected during no ice cover (2020).

To investigate how ice cover contributed to the ∼50% dissimilarity in winter diatom expression, DE analyses were performed. These results indicated 354 genes belonging to putative Bacillariophyta were differentially expressed (|Log_2_FC| ≥ 2, p_adj_ < 0.05), with 311 of these genes increasing in relative expression in ice free samples (variable of interest) and 43 decreasing (Supplemental Table 1X). Diatoms of the class Mediophyceae had the highest representation, comprising ∼50% of DE genes while other classes formed a net total of ∼10% (40% unclassified diatoms) (Supplemental Figure 13A). Further analysis revealed 33% of the polar centric DE genes were annotated as *Chaetoceros*-like (Supplemental Figure 13B), despite *Chaetoceros*-like genes forming ≤ 10% of mapped reads throughout the winter libraries (Supplemental Figure 7). Here, the “*Chaetoceros*-like” label arises as the transcriptomes were annotated with largely marine-comprised databases due to a lack of comprehensive freshwater taxonomic sequencing [9, 59]. Hence, diatom class are reported in-text, genera are reported in the Supplemental.

Genes categorized in COG category C (Energy production and conversion) were the second highest represented category within the DE dataset, with most genes exhibiting increased expression within ice-free diatom communities (Figure 5). Of these genes, 64% belonged to the class Mediophyceae (Figure 5A, Supplemental Figure 14). Notably, the expression of genes encoding for iron-containing photosynthetic proteins such (ferrodoxin-*petF*, flavoprotein-*etfA*, and ferritin-*ftnA*) increased in relative expression in ice-free communities while expression of photosystem II-*psbA* decreased (Figure 5B, C). Likewise, relative expression of genes within COG category P (Inorganic ion transport and metabolism) increased in ice-free samples (Supplemental Figure 15 A,), with expression of putative iron transporting genes (OMFeT_1-3) increasing in ice-free communities. DE genes within COG category P also largely belong to the Mediophyceae class, comprising ∼40% of the annotated genes (Supplemental Figure 15, 16). Notably, two proton-pumping rhodopsin genes (PPRs. COG S), which were most recently found to be a light-driven, retinal-based alternative to classical phototrophy in a cold-adapted freshwater photosynthetic bacterium [60], significantly increased in expression within ice-free diatom communities (Figure 6A, Supplemental Figure 17A). Further, the expression 9 additional diatom PPRs increased within the ice-free water column, though they fell short of differential expression (Supplemental Figure 17B). Taxonomic annotations demonstrated these 11 PPRs largely belonged to the Fragilariophyceae (33.33%) and Mediophyceae (22.22%) classes (Unclassified Diatoms = 44.44%), with the two DE PPRs annotated at the phylum (PPR_1, Bacillariophyta) and genus level (PPR_2, *Pseudo-nitzschia fraudulenta*-like). Phylogenetic analyses suggested diatoms horizontally acquired PPRs from bacteria, as there is evidence for at least 3 instances of horizontal gene transfer within our analysis (Figure 6C). The majority of the PPRs in our study clustered with or near Eukaryotic rhodopsins. Notably, the most highly DE PPR in our study (PPR_2, gene 538736) clustered closely with the marine diatom PPR belonging to *Psuedo niztschia granii* (Figure 6C) [61, 62]. Refer to Supplemental Methods/Results and Supplemental Tables for further details.

**Figure 5:**
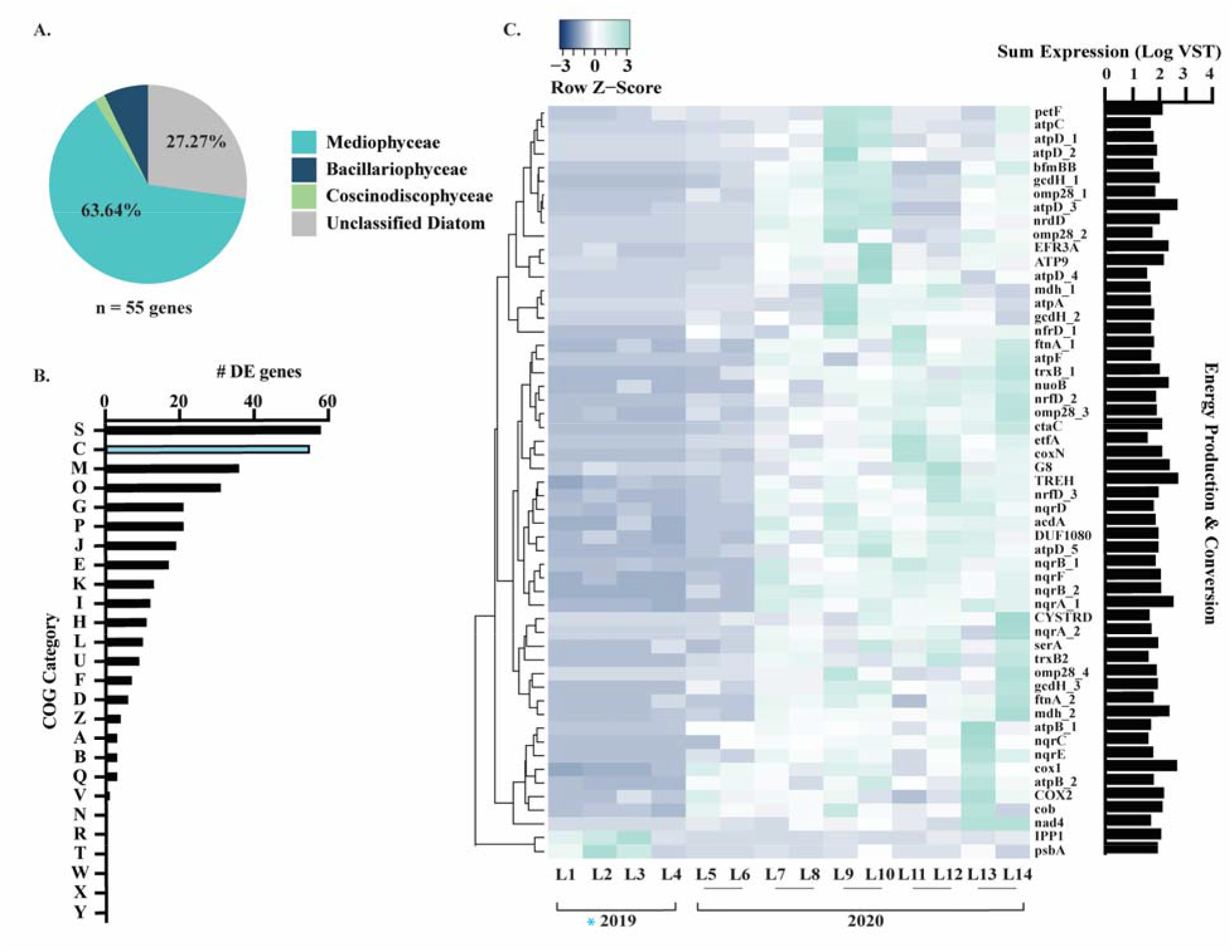
Bacillariophyta energy production and conversion transcript abundance patterns in response to ice cover. **A)** Taxonomic distribution of DE genes categorized within COG category C (Energy Production and Conversion). **(B)** COG assignments for all 354 DE genes in response to ice cover, with COG category C indicated in blue. **(C)** Heatmap depicting COG category C DE gene expression (VST) in response to ice cover across the 14 winter libraries.

**Figure 6:**
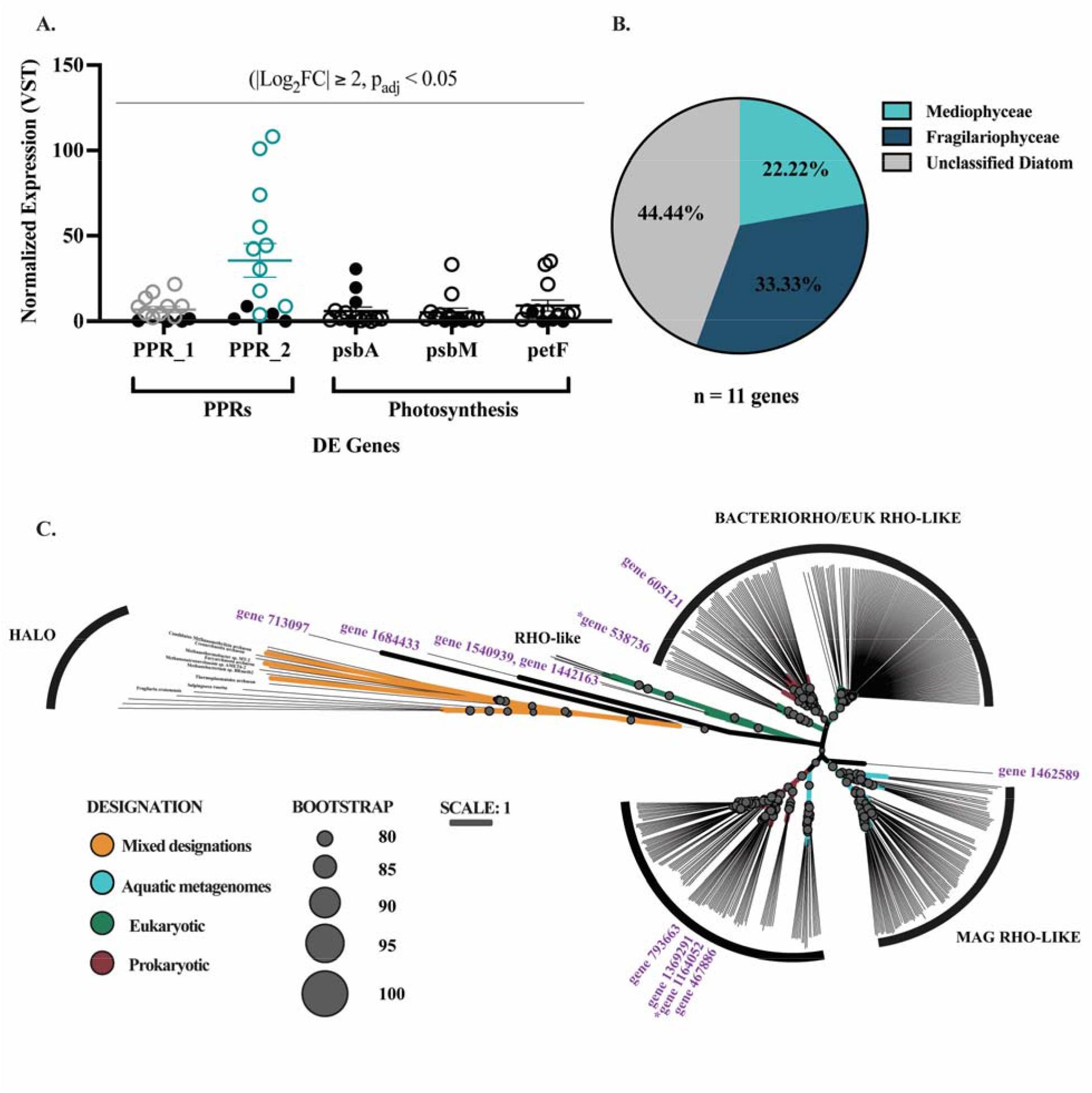
Bacillariophyta PPR transcript abundance patterns in response to ice. **(A)** Normalized expression (VST) of two DE genes functionally annotated as PPRs (PPR_1 (grey), PPR_2 (teal)) and representive DE genes functionally annotated to be involved in photosynthesis (black). Photosynthetic genes were selected because they are involved in light-harvesting (*psb*A, *psb*M) or the transfer of electrons along the photosystems (*pet*F). Each circle corresponds to gene expression in one of the 14 libraries. Solid black circles indicate the sample was collected during ice cover (2019) open shapes indicate the sample was collected during no ice cover (2020). **(B)** Taxonomic distribution of the 11 genes functionally annotated as PPRs (and confirmed with subsequent phylogenetic analysis). **(C)** Phylogenetic tree of PPR distribution within diatoms. Study putative rhodopsin-like proteins (n = 11, pink) were distributed within several rhodopsin groups and sub-groups to determine likelihood of putative genes being of bacterial or eukaryotic origins. The position of study genes is labelled by their associated /groups, with the exception of genes 713097 and 1684433 being GCPR transmembrane rhodopsin associated proteins, and gene 1462589 being unclear (most closely associated with the genes of metagenomic origin. Bootstrap values are based off 1000 replicates and are identified if above 80. Abbreviations: HALO; Halorhodopsin, XANTHO; xanthorhodopsin, METAGENOME RHO-LIKE; metagenomic origin rhodopsin-like putative proteins, BACTERIORHO/EUK RHO-LIKE; bacteriorhodopsins and eukaryotic origin rhodopsin-like putative proteins, RHO-like; sensory eukaryotic rhodopsin-like proteins. The two DE PPRs are indicated with asterisks.

DE analyses in response to season were performed with diatom libraries to identify trends truly unique to the ice cover DE dataset. The top 10 COG categories represented in each dataset overlapped except for COG category M, which was the third most abundant in ice cover analyses compared to the twelfth most abundant in season analyses (Supplemental Figure 18). Further analysis of these COG M (Cell wall, membrane, and envelope biogenesis) genes revealed 58% belonged to the class Mediophyceae (Figure 7A, Supplemental Figure 19). Intriguingly, 50% of the DE COG M genes encode for fasciclins (FAS1) which increased in expression under ice-free conditions (Figure 7B, C). Fasciclins are secreted glycoproteins involved in diatom cell-cell adhesion and cell-extracellular matrix adhesion [63, 64]. All 18 DE fasciclin genes were either assigned to the class Mediophyceae or unclassified beyond the phylum level (Bacillariophyta). Phylogenetic analyses indicated diatoms horizontally acquired FAS1 from bacteria, as there is evidence for at least 6 instances of horizontal gene transfer within our analysis (Figure 7D). Broadly, the FAS1 domain is widely distributed in diatoms, with ∼140 marine and freshwater diatoms found to contain this protein domain including the model cold-adapted diatom *Fragilariopsis cylindrus* [65]. Refer to Supplemental Methods/Results and Supplemental Tables for further details.

**Figure 7:**
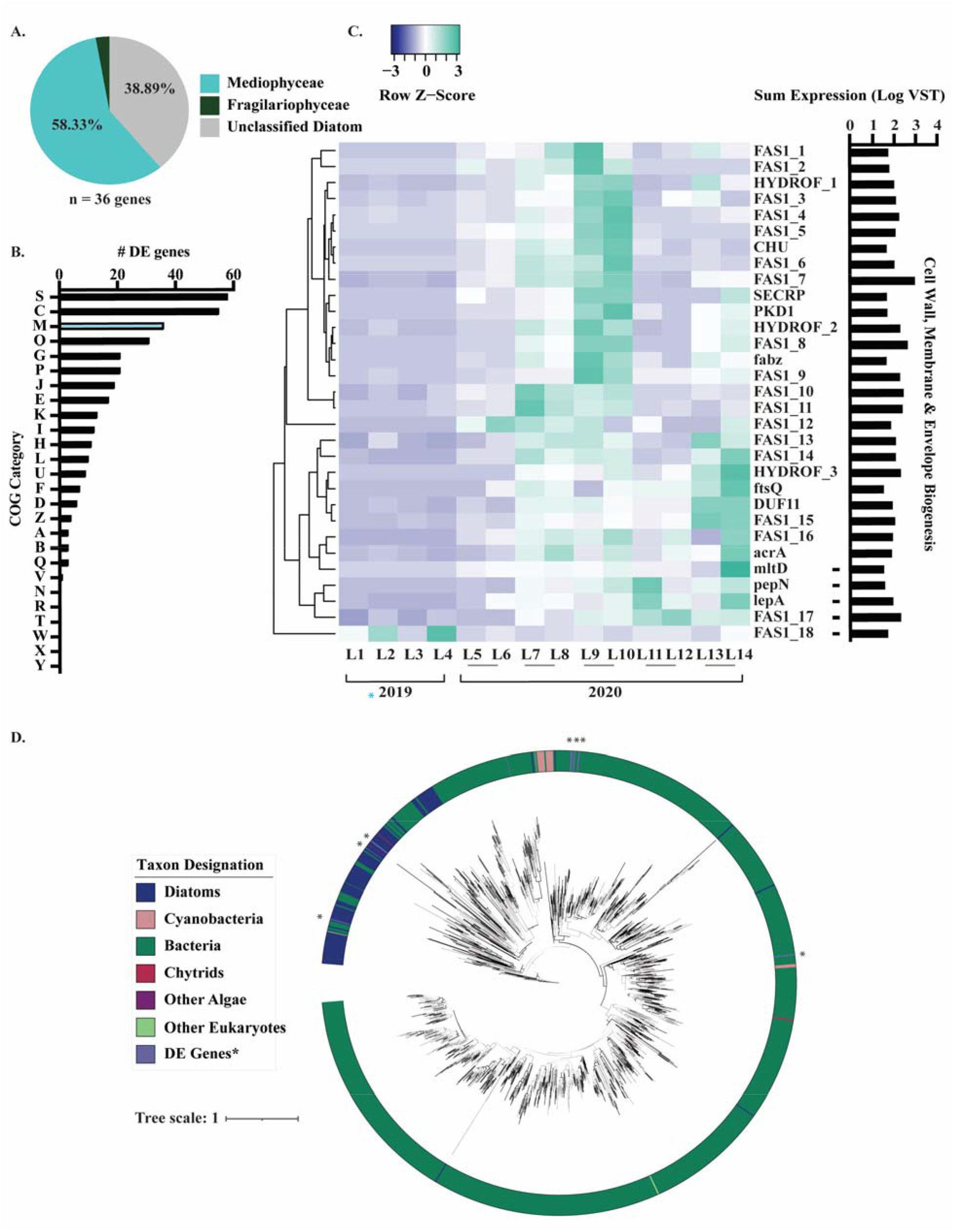
Bacillariophyta fasciclin transcript abundance patterns in response to ice cover. **(A)** Taxonomic distribution of DE genes categorized within COG category M (Cell wall, membrane, envelope biogenesis). **(B)** COG assignments for all 354 DE genes in response to ice cover, with COG category M indicated in blue. **(C)** Heatmap depicting COG category M DE gene expression (VST) in response to ice cover across the 14 winter libraries. **(D)** Phylogenetic tree of fasciclin distribution within diatoms. Bootstrap values above 70 are indicated with black lines. The FAS1 domain was found in 141 marine and freshwater diatoms of diverse ecological habitats (indicated in blue). The 18 DE diatom fasciclins in this study are indicated in purple with asterisks. Bacterial fasciclins indicated in dark green), cyanobacteria (pink), chytrids (red), other algae (purple), other eukaryotes (light green).

## Discussion

The present study examined how phytoplankton (specifically diatoms) are responding to rapidly declining ice cover in a northern temperate lake (Lake Erie). Indeed, ice cover has declined by ∼70% on the Laurentian Great Lakes over the 45-year period from 1973-2017 [66]. Notably, it has been suggested this decline in ice cover increases light limitation in shallow lakes such as Lake Erie, hence we investigated this hypothesis that diatoms are light limited in the turbid ice-free water column [4]. In this study, we demonstrated ice-free conditions decreased diatom bloom magnitude and altered composition. Notably, diatoms exhibited increased relative expression of photosynthesis and iron-transport genes under ice-free conditions: trends which are consistent with light limitation [67, 68]. Further, metatranscriptomic analysis provides evidence for two novel hypotheses concerning diatom adaptations to the ice free state of the lake: (1) *The ice cover PPR energy hypothesis* and (2) *The fasciclin mediated rafting hypothesis*. We provide this information couched within the context of the ecophysiological implications of a climatically altered future for psychrophilic aquatic communities.

### Ice free conditions alter diatom bloom magnitude and composition

It was previously noted *A. islandica* biovolumes were 95% decreased in the ice-free turbid water column (2012) compared to ice-covered conditions the years prior (2010, 2011); with light limitation cited as the potential driver of this trend [4]. By comparison, Chl *a* biomass, centric diatom counts, and *A. islandica* counts did not significantly decrease relative to ice cover in our study, though they all exhibited a consistent declining trend. However, our findings are in juxtaposition to a prior Lake Erie winter study (2007-2010) which reported abundances of *A. islandica* and *Stephanodiscus* spp. 1-2 magnitudes higher, with Chl *a* concentrations supporting this trend [3]. In addition, subsequent studies also reported diatom communities overwhelmingly dominated by *A. islandica,* with *Stephanodiscus spp.* present in lesser concentrations [3, 9–11, 69]. In contrast, we discovered cell abundances of *Stephanodiscus* spp. were significantly higher than *A. islandica* in the ice-covered community. We hypothesize a decrease in consecutive years of high ice cover may drive this decline of *A. islandica* dominance (Figure 1D). Regardless, our results indicate the winter diatom bloom community has markedly declined in magnitude and altered in composition since prior winter Lake Erie surveys (2007-2012).

In turn, it has been suggested that declines in diatom biomass may cause this niche to be filled by cryptophytes and dinoflagellates, as mixotrophs are suggested to be better suited for the turbid water column [4, 7]. While we observed significantly higher abundances of these groups in ice-free samples within our study, their cellular and transcriptional abundances remained below centric diatoms by an order of magnitude. Hence, our results demonstrate that low ice cover during this season did not induce significant large-scale phyla-level shifts in major eukaryotic phytoplankton community composition as previously suggested. Cumulatively, this implies future ice-free winter communities may remain dominated by centric diatoms as observed in this study, albeit at a lesser magnitude.

While net centric diatom abundances did not significantly differ by ice conditions in our study, diatom community composition exhibited significant changes at the genus level. Cell abundances of *Stephanodiscus* spp., were ∼50% lower in ice-free samples, resulting in ∼equal abundances of *Stephanodiscus* spp. and *A. islandica* within the ice-free water column. Further, small centric diatom taxa (5-20 μm size) were mainly absent in samples from ice-covered sites yet formed 10-82% of total diatom counts in ice-free sites, with a bloom of these taxa noted at site 8. This suggests ice-free conditions may increase populations of smaller, centric diatoms in future warmer and ice-free winters. The trend is supported by prior studies which demonstrate warming temperatures decrease phytoplankton cell size [70] and select for smaller taxa [71, 72]. We also noted significant increases in pennate diatom abundance in the ice-free water column. Cumulatively, if these observations represent long-term trends, future ice-free diatom communities will be more diverse with lower biomass.

### Evidence of light limitation within the ice-free water column

The relative expression of photosynthesis-associated genes increased overall within ice-free diatom communities, suggesting potential efforts to increase light capture and light-driven processes within the turbid water column. Most notably, we observed an increase in expression of iron transporters coinciding with various genes encoding for iron-rich photosynthetic structures and photosystem components. In support, a prior study found temperate phytoplankton acclimate to low light conditions by increasing their number of iron-rich photosystems [68]. Thus, our data suggest freshwater diatoms in the ice-free, turbid Lake Erie water column were attempting to build additional photosystems in response to decreased light availability. Indeed, our data offers transcriptional support for a prior study which found primary production rates (measured via C^14^ tracing) to be lower within the Lake Erie ice-free water column (2012) compared to the ice covered water column (2010-2011) [4]. Further, two diatom PPRs within our dataset increased within the ice-free water column while *psb*A coincidingly decreased, suggesting diatoms may be attempting to use alternative phototrophic strategies other than classical photosynthesis within the ice-free water column. Cumulatively, this suggests diatoms may be attempting to use alternative light-driven energy mechanisms as a means to evade classical light limitation. Other studies demonstrate enlarged light antennae is another response to light limitation, as this increases light harvesting [73] and is suggested to be particularly advantageous in cold environments [67]. While we did not observe evidence of this phenomenon in our dataset, this may have occurred prior or post-sampling of the community as metatranscriptomics offers only a “snapshot” episodic glimpse at community response. Hence, while we found supportive evidence of light limitation in this study, further research is required in seasonally cold temperate systems.

### The role of PPRs as a function of ice cover

We observed increases in the expression of genes encoding for PPRs within the ice-free diatom community. PPRs are light-harvesting retinal-containing proton pumps distinct from the chlorophyll-containing antenna of classical photosynthesis [74], yet capable of absorbing as much light energy as Chl *a* [75]. On a global scale, it is thought microbial rhodopsin driven phototrophy is a major marine light harvesting process [76]. Beyond prokaryotes, they have been characterized within a number of marine diatoms [62] and dinoflagellates [77]. Notably, PPRs have been suggested to serve as an alternative light-driven energy source for marine diatoms under conditions which limit classical photosynthesis [62, 78]. However, compared to the marine literature [75, 79–83], PPRs are widely understudied in fresh waters. To our knowledge, this study is the first to report the presence of PPRs within freshwater diatoms to datalthough we are not the first to suggest PPRs may play a role in energy generation beneath ice. It was recently found that a photoheterotrophic bacterium isolated from an alpine lake used PPRs as an alternative phototrophy mechanism [60]. The authors of this study further hypothesized the contribution of PPRs to energy generation is linked to ice cover. Here, we present evidence diatoms within the ice-free Lake Erie water column increase the expression of PPRs in the ice-free, low-light turbid environment, which lends support to this hypothesis regarding an ecophysiological role of PPRs within icy freshwater environments.

Nonetheless, there are a variety of ecophysiological explanations for this PPR phenomenon. This may be an attempt to scavenge wavelengths of light beyond those absorbed by Chl *a*; lending further evidence they are light limited. Indeed, PPRs absorb at a maximum wavelength of ∼525nm (green light) [83] in contrast to Chl *a* which absorbs at a 430-470nm (blue) and 660-670nm (red). Hence, PPRs would allow diatoms to access alterative light niches in the turbid water column. This is further supported in our data, as expression of two diatom PPRs was significantly higher in the ice-free water column whereas the expression of *psb*A was significantly higher in ice-covered samples (Figure 6A). Alternatively, green light penetrates up to 100m depth, with only to blue light penetrating further. Hence, PPRs may be involved in light-acquisition during well-mixed isothermal conditions when diatoms would be mixing throughout the benthic and pelagic environments in shallow lakes (Supplemental Figure 20). Broadly, this suggests an ecophysiological role of PPRs for diatoms across global freshwater and marine environments. Thus, our observations suggest that the role of these proton-pumping rhodopsins within fresh waters demands more attention [84], as it is possible that in future scenarios (less ice cover, more turbidity) they may serve as important evolutionary selectors.

### The role of fasciclins in the ice-free turbid water column

Though fasciclins (FAS1) remain widely uncharacterized in diatoms, prior studies have described fasciclins within the diatom species *Amphora coffeaeformis* [64] and *Phaeodactylum tricornutum* [63]. Both studies identified fasciclin proteins within diatom-secreted exopolymer substance adhesion trails and concluded these molecules facilitate diatom motility, adhesion, and aggregation. In this study, 58% of the DE diatom fasciclins belonged to the class Mediophyceae (27% to unclassified diatoms). As a result, we hypothesize Mediophyceae diatoms were rafting *via* cell-adhesion fasciclins to optimize their location within the ice-free, turbid Lake Erie water column, thus avoiding light limitation.

This hypothesis is largely based on a similar “rafting” strategy that is well-documented in centric marine diatoms, notably *Rhizosolenia* spp. [85–88].These studies demonstrated rafts become positively buoyant in response to a variety of physiological stressors, including iron limitation [86, 89, 90]. Further, our phylogenetic analyses demonstrated this fasciclin-mediated rafting hypothesis may not be unique to Lake Erie diatoms alone. Fasciclins were identified in ∼140 marine and freshwater diatoms including the model polar marine diatom *Fragilariopsis cylindrus* [91]. Hence, this further implies an ecophysiological role exists for these proteins in the globally frigid waters. In turn, this also indicates further research is required regarding the role of fasciclins within polar aquatic systems and psychrophilic organisms broadly, especially when considering the rapid global decline in ice cover.

## Conclusions

Lakes are sentinels of climate change [92]. Indeed, our study builds on data which demonstrate community-wide responses to climate-driven declines in ice cover are already underway. Notably, we provide evidence which suggests diatom declines are driven by light-limitation in the turbid ice-free water column (Figure8). Indeed, Ozersky et al., [7] suggested warmer winters will induce a change in the Great Lakes mixing regime, shifting from dimictic mixing patterns to a warm monomictic mixing pattern characterized by continuous isothermal conditions throughout winter. Hence, adaptations to evade coinciding exacerbations in light limitation (such as the possession of PPRs and FAS1 described in this study) may be of increased importance in future winter diatom survival as phytoplankton adapt to ice-free winters. Indeed, our data suggests climate change may not be just a “temperature” problem in the case of shallow temperate lakes, but a “light” problem. Regardless, with diatoms previously described as “one of the most rapidly evolving eukaryotic taxa on Earth” [93, 94] and prone to promiscuous horizontal gene transfer events [95], it would be surprising if they failed to adapt to an ice-free future. Ultimately, we cannot place the consequences of the metatranscriptomic observations we describe in a quantitative framework. To this end, our observations, which demonstrate variability associated with conditions consistent with projected future climate scenarios, carve out a critical path forward and provide cautionary insight of what may be yet to come in global temperate lakes.

**Figure 8:**
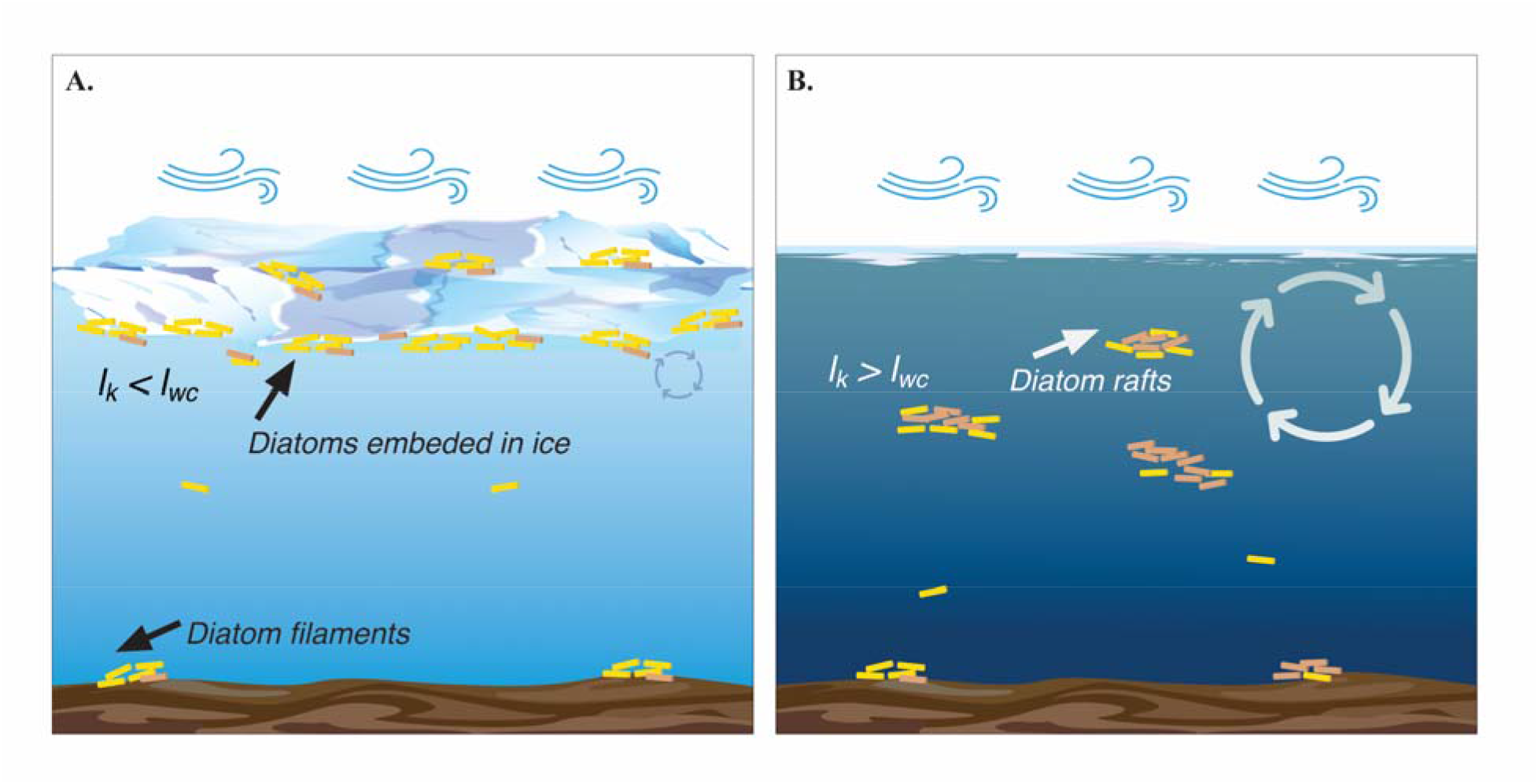
Schematic representing how ice cover alters freshwater diatom colocation strategy throughout the water column. **(A)** Ice covered water column which exhibits minimal convective mixing. As a result, I_k_ < I_wc_ (where I_k_ = irradiance at which photosynthesis is light saturated and I_wc_ = mean water column irradiance). Iwc is calculated based on light extinction coefficient and mixing depth – which in an ice-free winter (i.e., holomictic state) is the bottom, while in the presence of ice-cover, is limited to shallow convective mixing. Diatoms nucleate ice and partition to the surface ice cover within the photic zone. **(B)** Ice-free water column which exhibits isothermal conditions and thorough mixing. As a result, I_k_ > I_wc_, and light-limited diatoms express fasciclins to forms rafts of increased buoyancy to optimize their ability to harvest light in the turbid water column. Beige colored diatoms in both diagrams represent diatoms that possess PPRs, with an increased number of PPR-possessing diatoms selected for in the ice-free turbid water column in panel B.

## Data availability

Raw and processed reads for the data used in this study are available through the JGI Data Portal (https://data.jgi.doe.gov) under Proposal ID 503851.

## Author Contributions

This project was designed by SW and RMLM. Samples were collected by RMLM, TF, GB, and JC. Phytoplankton enumeration and identification was performed by WC. RNA extractions in addition to quality/quantity assessment were performed by BZ. Metatranscriptomic processing was performed by BZ, NG, EC and LD using a pipeline established by NG. Python scripts associated with metatranscriptomic pipeline were written by AT. The phylogenetic tree and corresponding analyses were performed by EC. Statistical analyses were made by BZ. Figures and the first full draft were made by BZ. All authors contributed to the revisions and final version of the manuscript.

## Acknowledgements

We are grateful to the command and members of the U.S. Coast Guard Cutter Neah Bay, Canadian Coast Guard Ship *Limnos* and M/V *Orange Apex* for their help with sample collection and the generation of hydrochemistry data. We thank Daniel H. Peck, James T. Anderson, Derek Niles and Arthur Zastepa for their help with sample collection and pre-processing. We thank Christa Pennacchio with the JGI for her coordination and sequencing expertise. We thank Dr. Gary LeCleir, Katelyn Houghton, Dr. Erik Zinser, and Dr. Jill Mikucki for their comments and suggestions. This work was funded by the National Institutes of Health, NIEHS grant 1P01ES023-28939-01, National Science Foundation grant OCE-1840715 (GB, RMLM, JC, SW), NSERC grant RGPN-2019-03943 (RMLM) and funding from the NSF GRFP DGE-19389092 (BNZ). This work conducted by the U.S. Department of Energy Joint Genome Institute (https://ror.org/04xm1d337), a DOE Office of Science User Facility, is supported by the Office of Science of the U.S. Department of Energy operated under Contract No. DE-AC02-05CH11231.”

## Additional Information

**Supplementary information** is electronically available at XXXXX.

**Correspondence** and requests for materials should be addressed to Steven W. Wilhelm or R. Michael L. McKay.

**Reprints and permission information** is available at http://www.nature.com/reprints.

