## Supplemental Figures for "Declines in ice cover induce light limitation in freshwater diatoms"

For publication in conjunction with the following:

*__Correspondence:__

| Library | Site | Ice cover | Season | Julian day | Month | Year |
|---|---|---|---|---|---|---|
| L1 | S1 | Yes | Winter | 57 | February | 2019* |
| L2 | S2 | Yes | Winter | 57 | February | 2019* |
| L3 | S3 | Yes | Winter | 70 | March | 2019* |
| L4 | S4 | Yes | Winter | 70 | March | 2019* |
| L5, L6 | S5 | No | Winter | 45 | February | 2020 |
| L7, L8 | S6 | No | Winter | 45 | February | 2020 |
| L9, L10 | S7 | No | Winter | 45 | February | 2020 |
| L11, L12 | S8 | No | Winter | 62 | March | 2020 |
| L13, L14 | S9 | No | Winter | 62 | March | 2020 |
| L15, L16, L17 | S10 | No | Spring | 122 | May | 2020 |
| L18 | S11 | No | Spring | 143 | May | 2020 |
| L19, L20 | S12 | No | Spring | 160 | June | 2020 |

**Supplemental Table 1. Metatranscriptomic libraries with respect to spatial, temporal, and climatic variables.** Libraries listed within the same row are biological replicates. Sample sites are numbered by chronological order of sampling (S1-12). Samples that were collected from under the ice are denoted with "Yes" and samples collected during a no ice cover are denoted with "No". Season is listed as either "Winter" or "Spring". Sample day is presented in Julian day, to account for interannual variability. Samples collected during 2019, a year of high ice cover overall throughout Lake Erie, are indicated with an asterisk. Samples collected during 2020, a year of low ice cover throughout Lake Erie, do not have an asterisk.

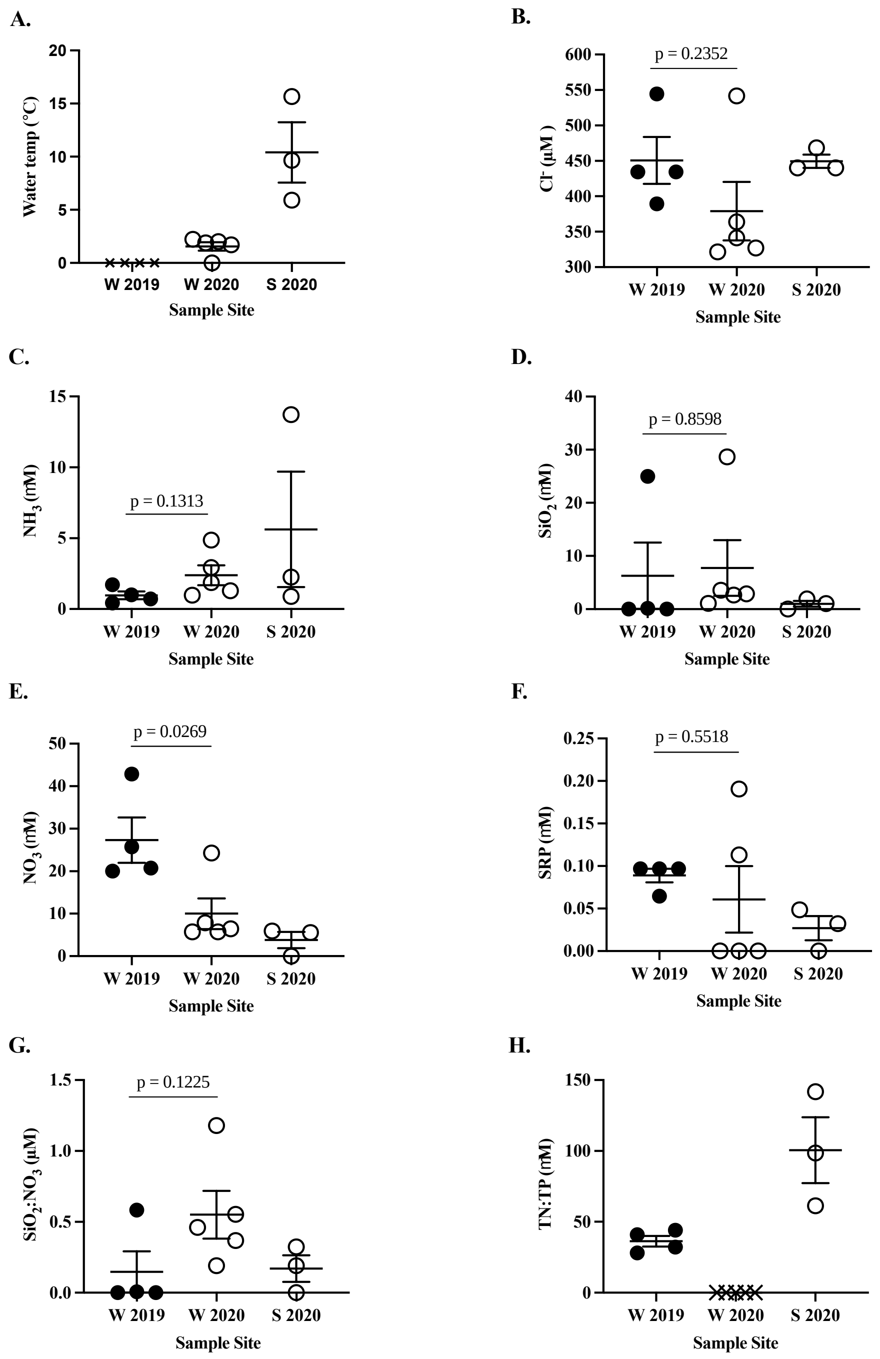


**Supplemental Figure 1:** Temperature and nutrient profiles (μM) across the 12 sample sites organized by season. W = winter (February-March), S = spring (May-June). Solid shapes indicate the sample was collected during ice cover (2019), open shapes indicate the sample was collected during no ice cover (2020**).** X's indicate there was no data reported. **(A)** Water temperature. **(B)** Dissolved Chlorine. **(C)** Dissolved Ammonia. **(D)** Dissolved Silicate. **(E)** Dissolved Nitrate. **(F)** Dissolved Soluble Reactive Phosphorus **(G)** Silica: Nitrate ratios. **(H)** Total Particulate Nitrogen: Total Particulate Phosphorus. Statistical comparisons as a function of ice cover were made using unpaired, two-tailed t-tests.

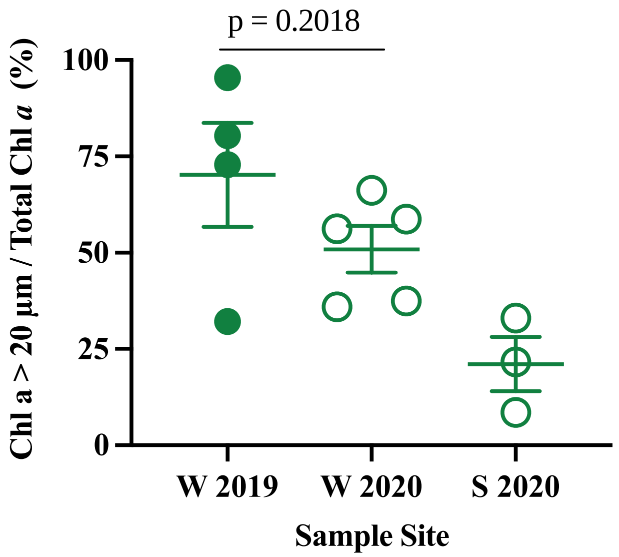


**Supplemental Figure 2:** Mean percent contribution of Chl *a* >20 µm to total Chl *a* across sample sites organized by season. W = winter (February-March), S = spring (May-June). Solid shapes indicate the sample was collected during ice cover (2019) open shapes indicate the sample was collected during no ice cover (2020). Statistical comparisons as a function of ice cover were made using unpaired, two-tailed t-tests.

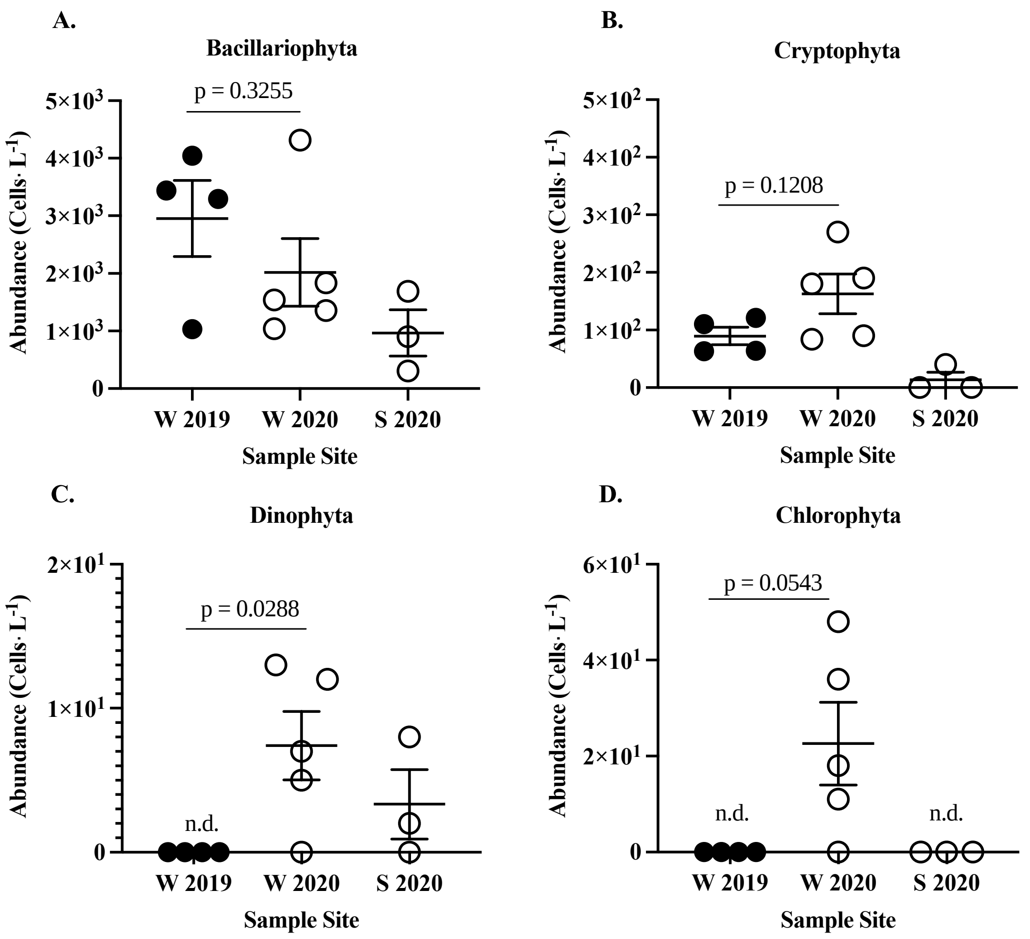


**Supplemental Figure 3:** Abundance (Cells•$L^{-1}$) of major eukaryotic phytoplankton taxa across sample sites organized by season. W = winter (February-March), S = spring (May-June). Solid shapes indicate the sample was collected during ice cover (2019) open shapes indicate the sample was collected during no ice cover (2020**).** Sites where the taxon was not detected are indicated with “n.d.”. **(A)** Sum diatom counts (*A. islandica* + *Stephanodiscus* spp. + centric diatoms of 5-20 µm + *Fragilaria* spp. + *Asterionella formosa*. + *Nitzschia* spp.). **(B)** Abundance of cryptophytes across sample sites. **(C)** Abundance of dinoflagellates across sample sites. **(D)** Abundance of chlorophytes. Statistical comparisons as a function of ice cover were made using unpaired, two-tailed t-tests.

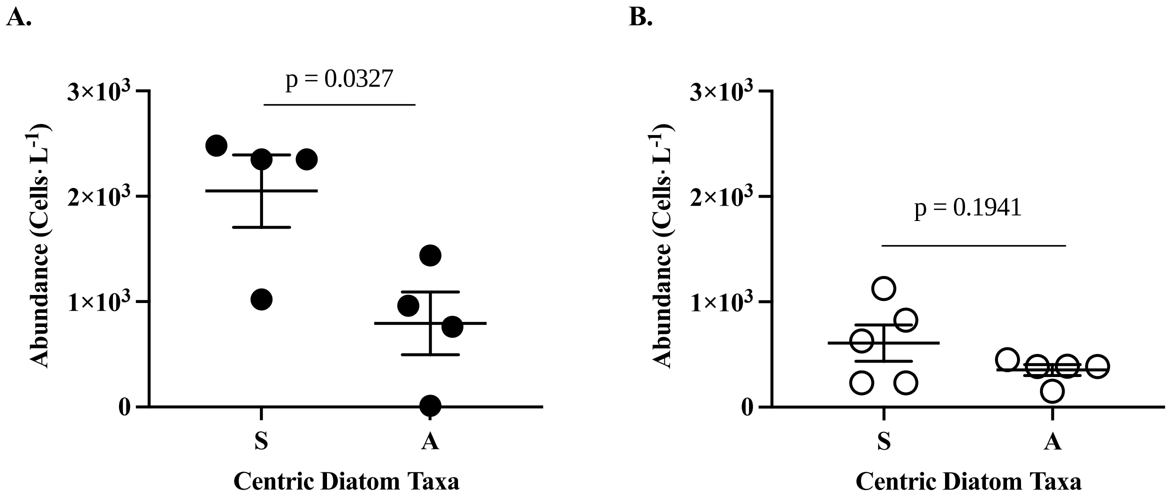


**Supplemental Figure 4:** Abundance (Cells•L$^{-1}$) of major centric, filamentous bloom-forming diatom taxa (S = *Stephanodiscus* spp., A = *A. islandica*) across winter sample sites organized by ice cover. Solid shapes indicate the sample was collected during ice cover (2019) open shapes indicate the sample was collected during no ice cover (2020**). (A)** Abundances of *Stephanodiscus* spp and *A. islandica* in winter ice-cover samples. **(B)** Abundances of *Stephanodiscus* spp and *A. islandica* in winter ice-free samples. Statistical comparisons as a function of ice cover were made using unpaired, two-tailed t-tests.

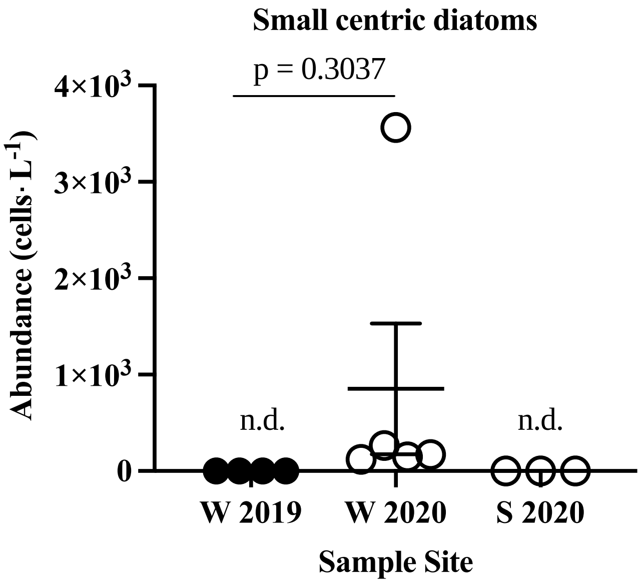


**Supplemental Figure 5:** Abundance (Cells•$L^{-1}$) of small centric diatom taxa (5-20 µm size) across sample sites organized by season. W = winter (February-March), S = spring (May-June). Solid shapes indicate the sample was collected during ice cover (2019) open shapes indicate the sample was collected during no ice cover (2020). Sites where the taxon was not detected are indicated with "n.d.". Statistical comparisons as a function of ice cover were made using unpaired, two-tailed t-tests.

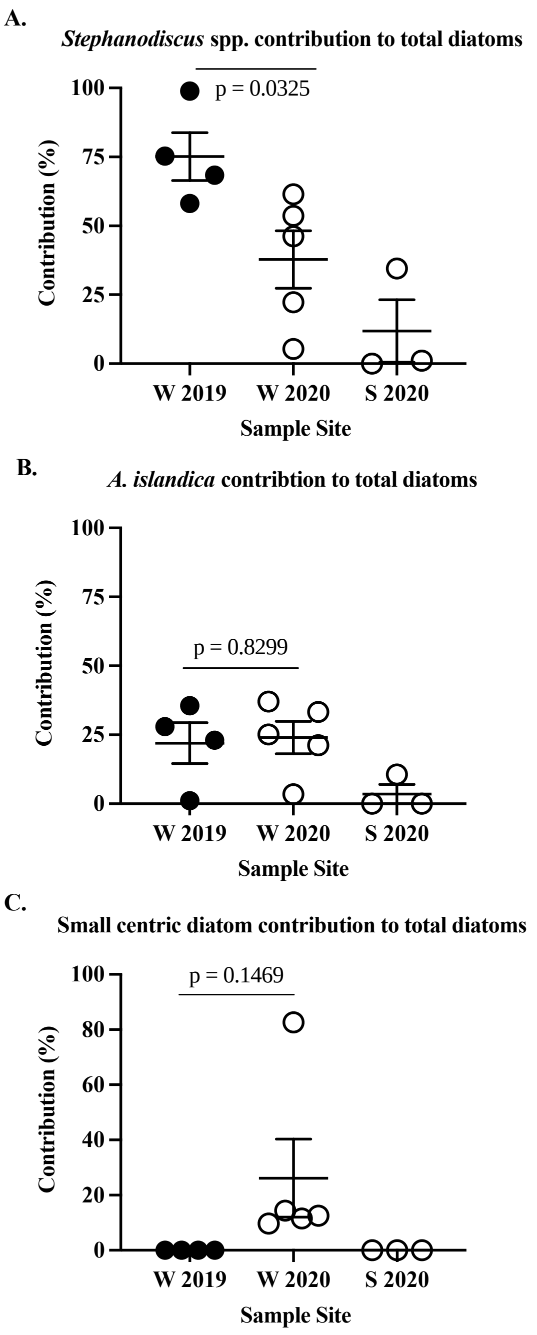


**Supplemental Figure 6:** Mean percent contribution of three centric diatom abundances to total diatom abundance across sample sites organized by season. W = winter (February-March), S = spring (May-June). Solid shapes indicate the sample was collected during ice cover (2019) open shapes indicate the sample was collected during no ice cover (2020). **(A)** Percent contribution of *Stephanodiscus* spp. genera abundance to total diatom abundance**. (B)** Percent contribution of *A. Islandica* abundance to total diatom abundance. **(C)** Percent contribution of small centric diatom taxa (5-20 μm size) to total diatom abundance. Statistical comparisons as a function of ice cover were made using unpaired, two-tailed t-tests.

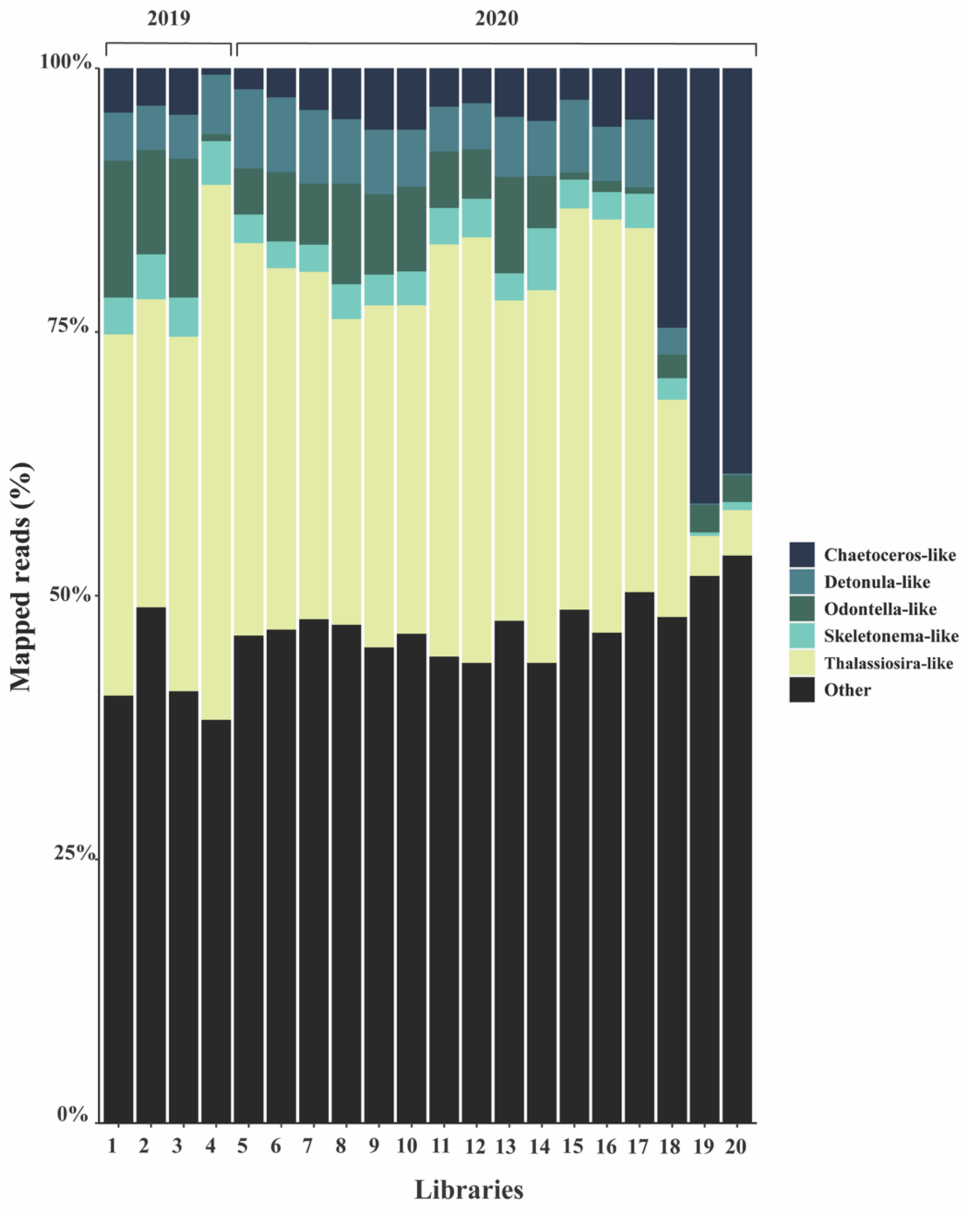


**Supplemental Figure 7:** Relative transcript abundance of Mediophyceae genera across the 20 libraries (listed in chronological order of sample date on the x-axis). All groups which formed <5% of the total mapped reads are included within "Other" (*Attheya* spp., *Cyclotella* spp., *Ditylum* spp., *Eucampia* spp., *Extubocellulus* spp., *Helicotheca* spp., *Minutocellus* spp., *Triceratium* spp). At the genus level, unclassified diatom counts were not included within total % read mapping calculations due to low-resolution at the genus level. They have been included within "Other" despite accounting for 59.04% of the total mapped reads to emphasize trends within classified genera.

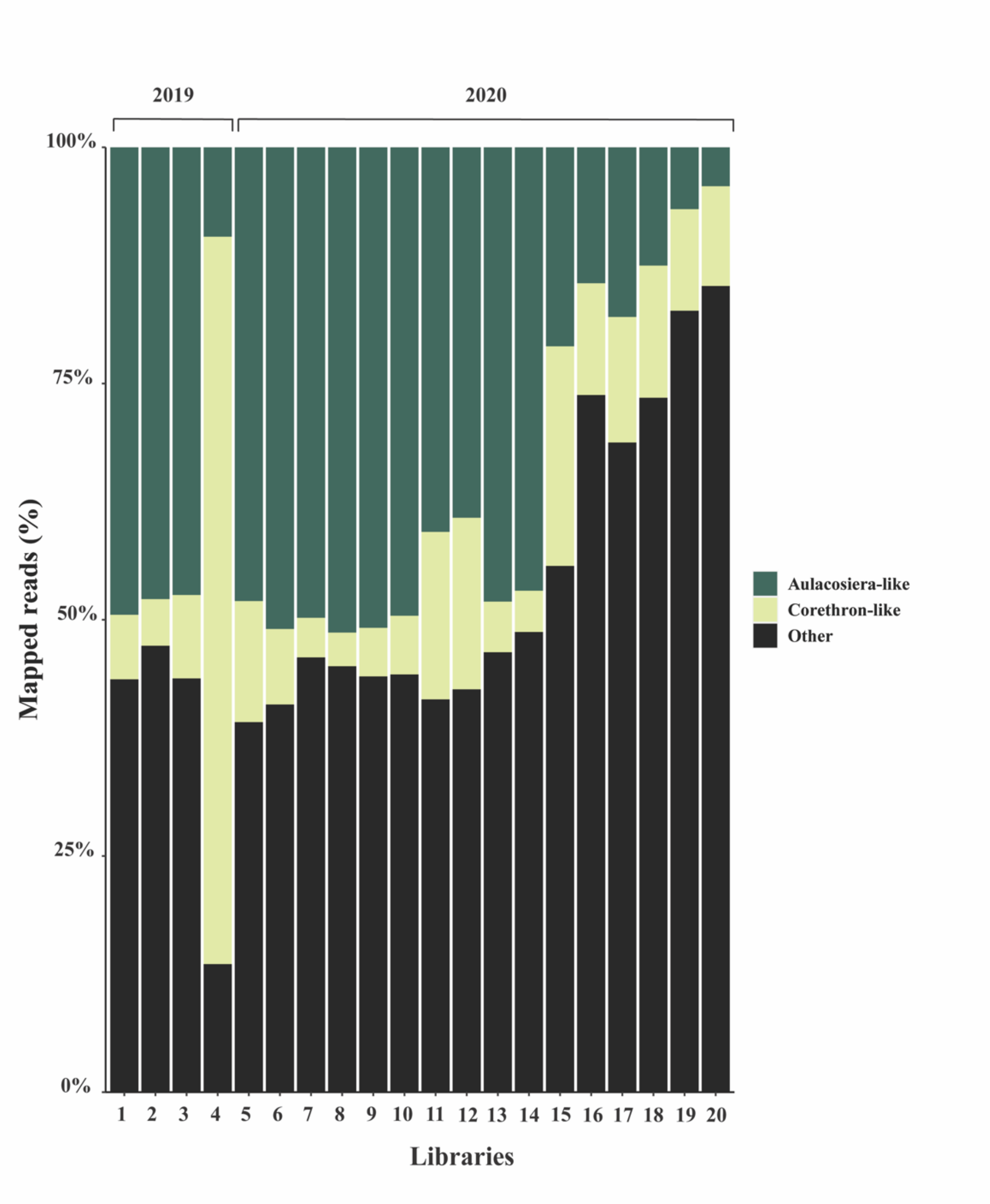


**Supplemental Figure 8:** Relative transcript abundance of Coscinodiscophyceae genera across the 20 libraries (listed in chronological order of sample date on the x-axis). All groups which formed <5% of the total mapped reads are included within "Other" (*Coscinodiscus* spp., *Dactyliosolen* spp., *Lepticylindrus* spp., *Proboscia* spp., *Rhizosolenia* spp., *Stephanopyxis* spp). At the genus level, unclassified diatom counts were not included within total % read mapping calculations due to low-resolution at the genus level. They have been included within "Other" despite accounting for 40.98% of the total mapped reads to emphasize trends within classified genera.

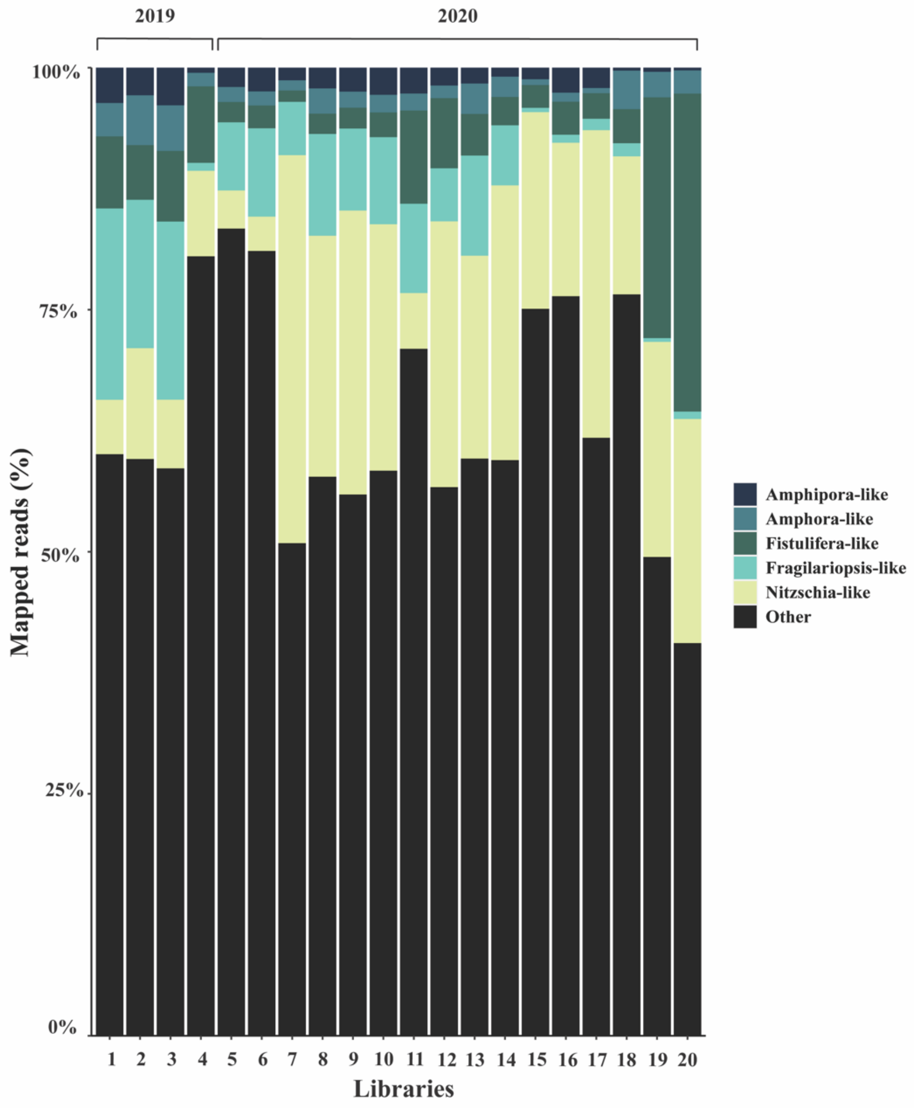


**Supplemental Figure 9:** Relative transcript abundance of Bacillariophyceae genera across the 20 libraries (listed in chronological order of sample date on the x-axis). All groups which formed <5% of the total mapped reads are included within "Other" (*Craspedostauris* spp., *Cylindrotheca* spp., *Entomoneis* spp., *Phaeodactylum* spp., *Psuedo-nitzschia* spp., *Stauroneis* spp). At the genus level, unclassified diatom counts were not included within total % read mapping calculations due to low-resolution at the genus level. They have been included within "Other" despite accounting for 66.00% of the total mapped reads to emphasize trends within classified genera.

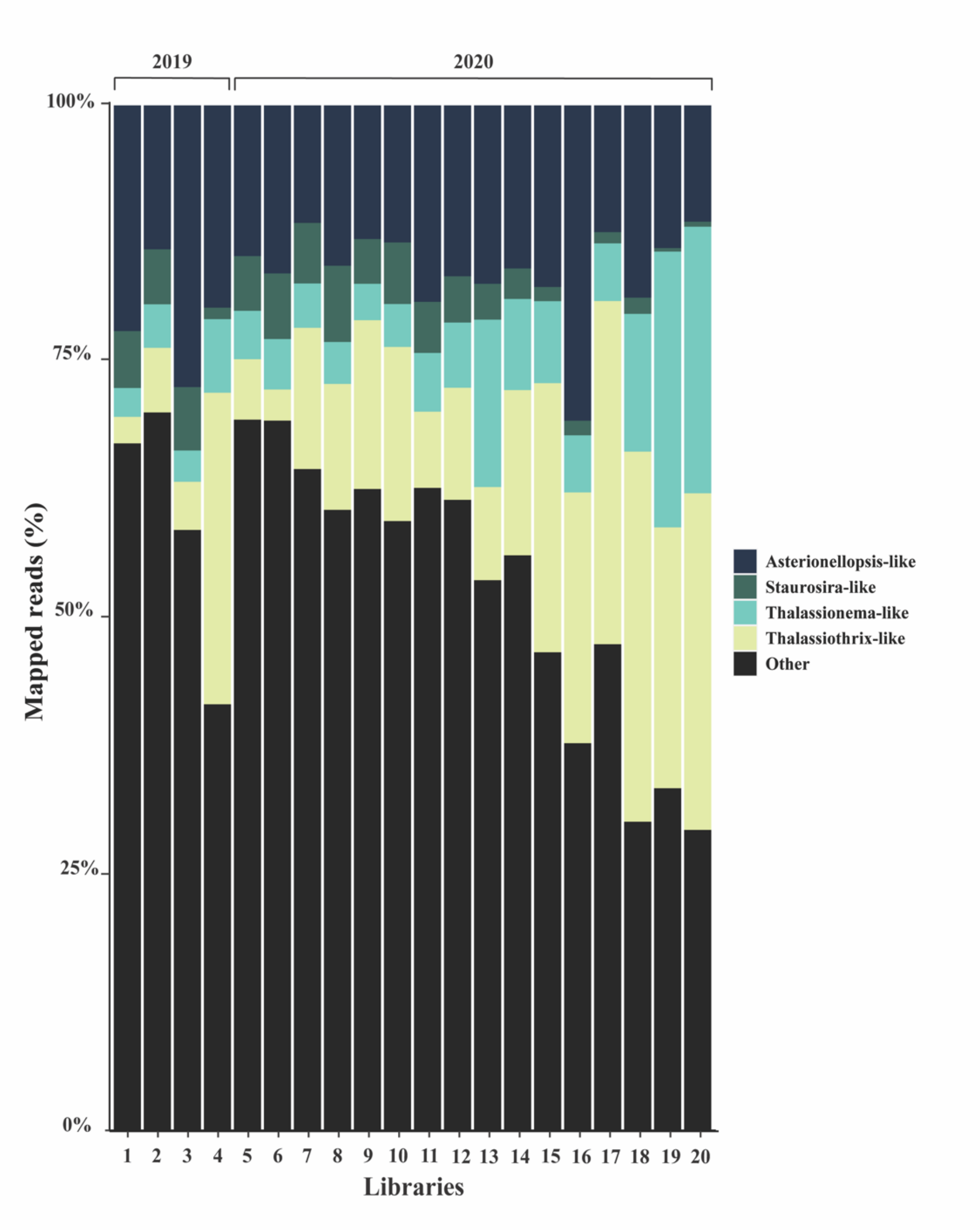


**Supplemental Figure 10:** Relative transcript abundance of Fragilariophyceae genera across the 20 libraries (listed in chronological order of sample date on the x-axis). All groups which formed <5% of the total mapped reads are included within "Other" (*Astrosyne* spp., *Cyclophora* spp., *Grammatophora* spp., *Licmophora* spp., *Striatella* spp., and *Syndropsis* spp.). At the genus level, unclassified diatom counts were not included within total % read mapping calculations due to low-resolution at the genus level. They have been included within "Other" despite accounting for 50.27% of the total mapped reads to emphasize trends within classified genera.

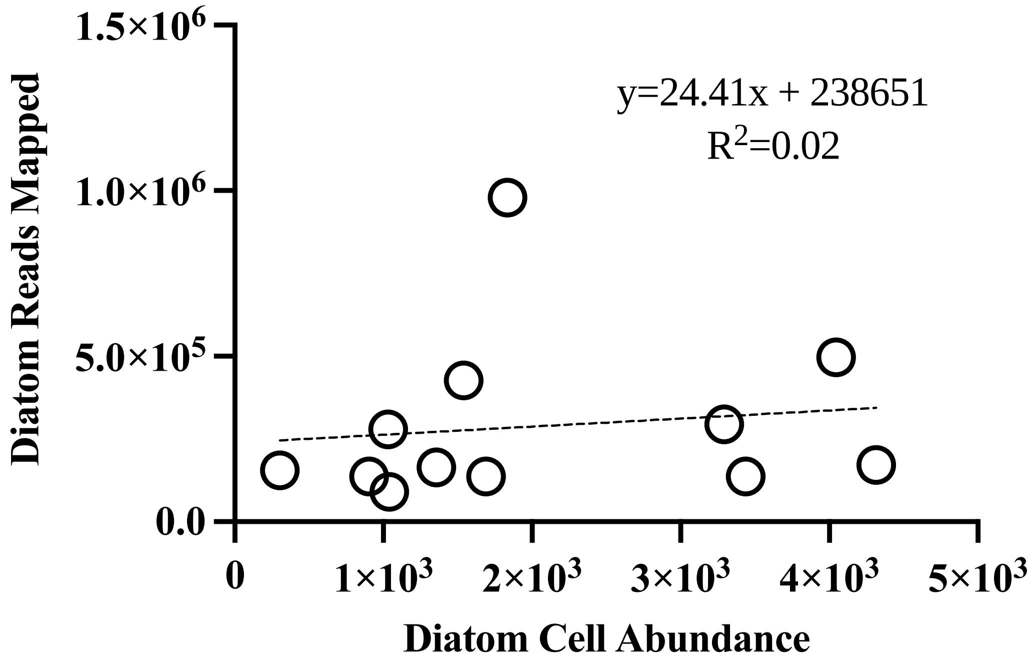


**Supplemental Figure 11:** Simple linear regression of diatom transcriptional abundances and diatom cell abundances.

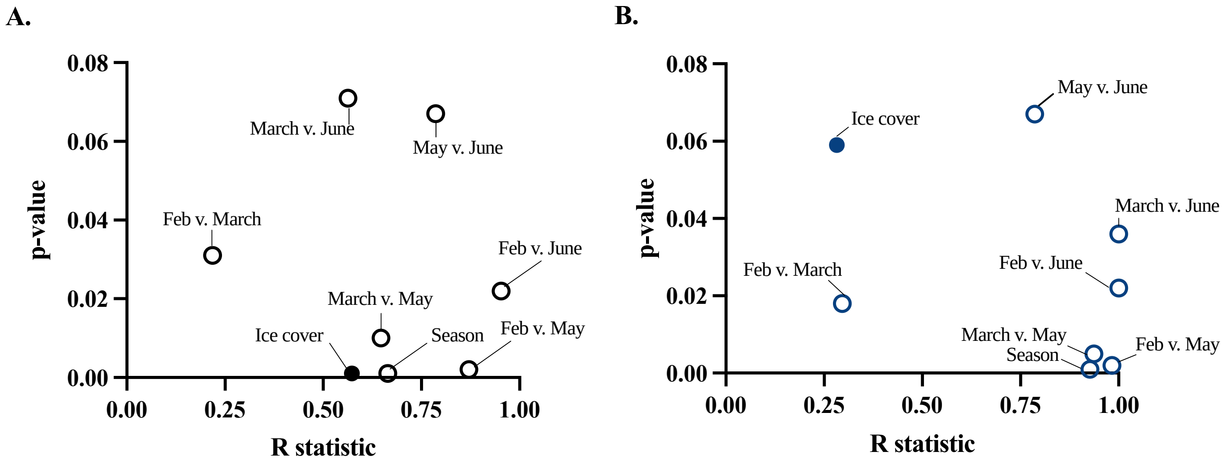


**Supplemental Figure 12:** ANOSIM tests plotted by R statistic and p-value. An R statistic close to 1.00 and a p-value < 0.05 indicates there is a significant difference between the 2 variables. Ice cover is indicated by a filled circle. (**A)** ANOSIM tests on the relative expression profiles (TPM) of the whole winter community comparing months, season (winter vs. spring), and ice cover (ice cover vs. no ice cover). (**B)** ANOSIM tests on the relative expression profiles (TPM) of the Bacillariophyta community comparing months, season (winter vs. spring), and ice cover (ice cover vs. no ice cover).

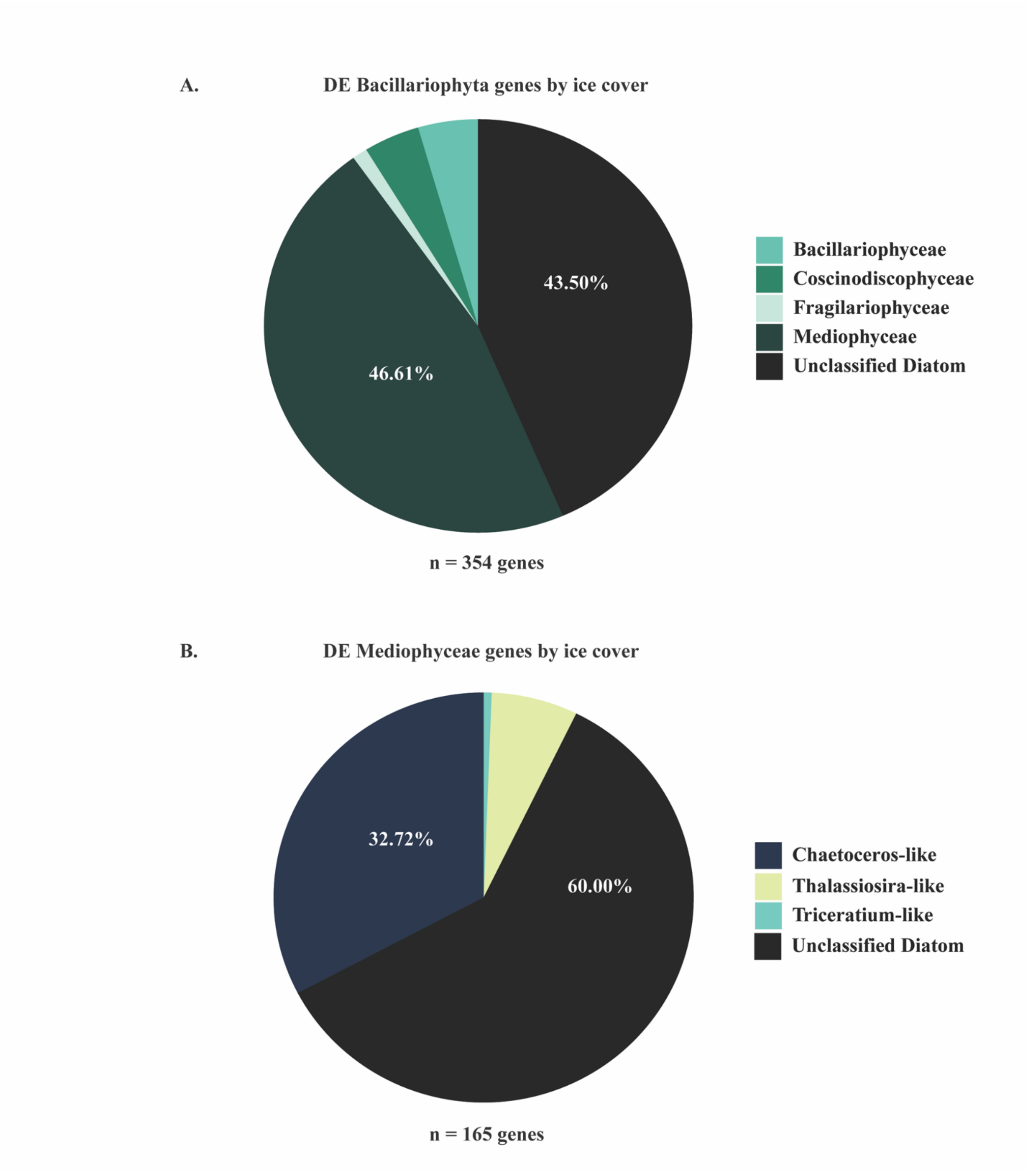


**Supplemental Figure 13:** Taxonomic distributions of diatom genes that were differentially expressed by ice cover. **(A)** Taxonomic distribution of all 354 DE genes by diatom class. **(B)** Taxonomic distribution of DE genes belonging to the Mediophyceae class.

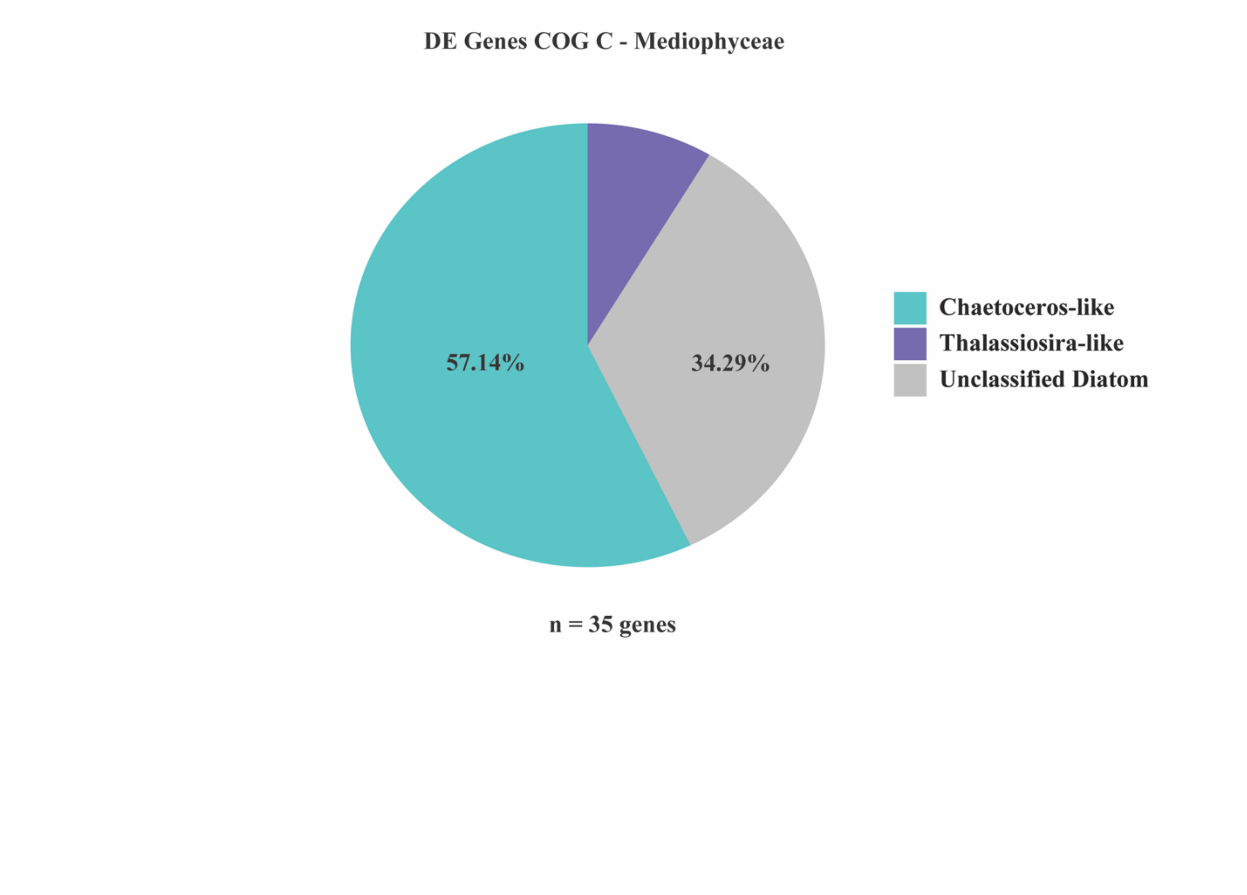


**Supplemental Figure 14:** Taxonomic distributions of Mediophyceae diatom genes that were differentially expressed by ice cover within COG Category C.

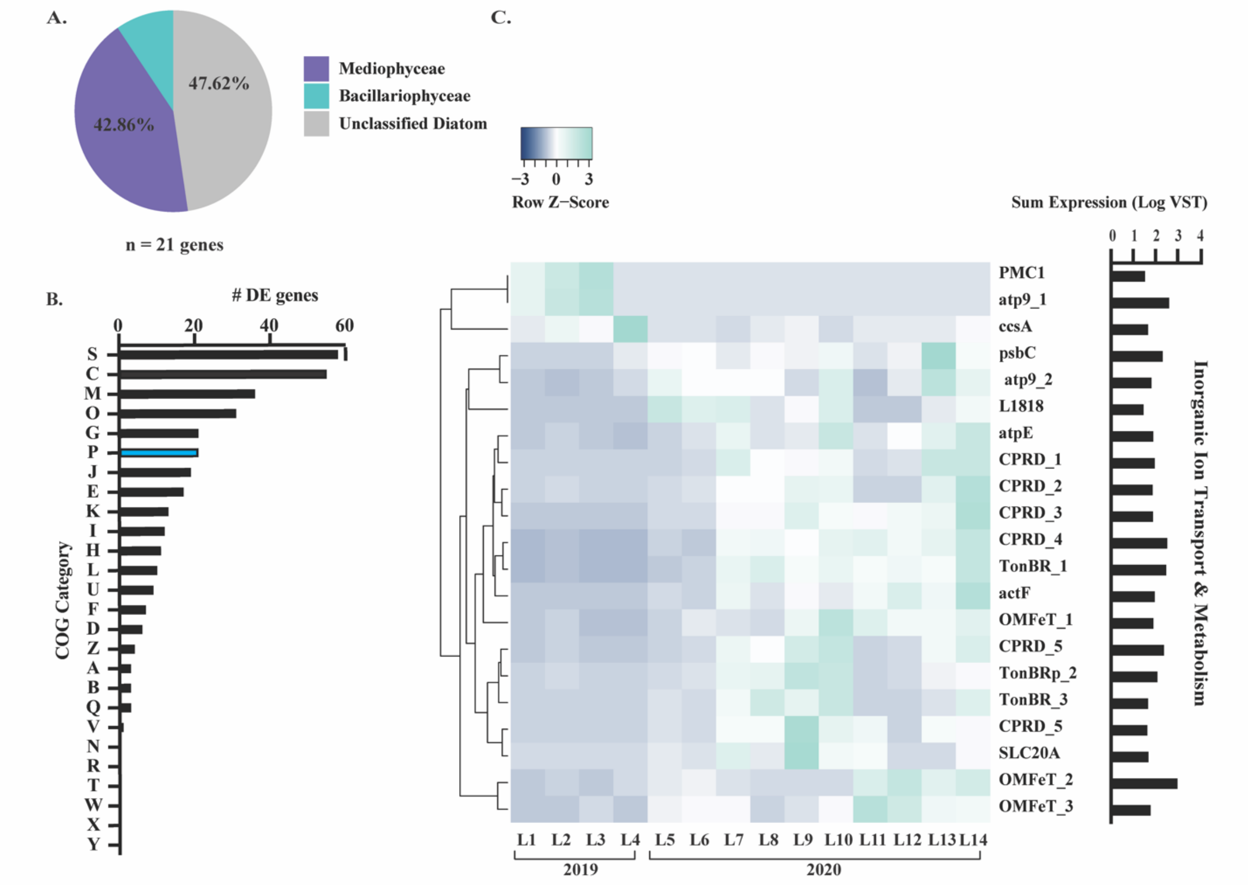


**Supplemental Figure 15:** Bacillariophyta transcript abundance patterns in response to ice cover-COG P (Inorganic ion transport and metabolism). **(A)** Taxonomic distribution of DE genes categorized within COG category P. **(B)** COG assignments for all 354 DE genes in response to ice cover, with COG category P indicated in blue. **(C)** Heatmap depicting COG category P differentially expressed gene expression (VST) in response to ice cover across the 14 winter libraries.

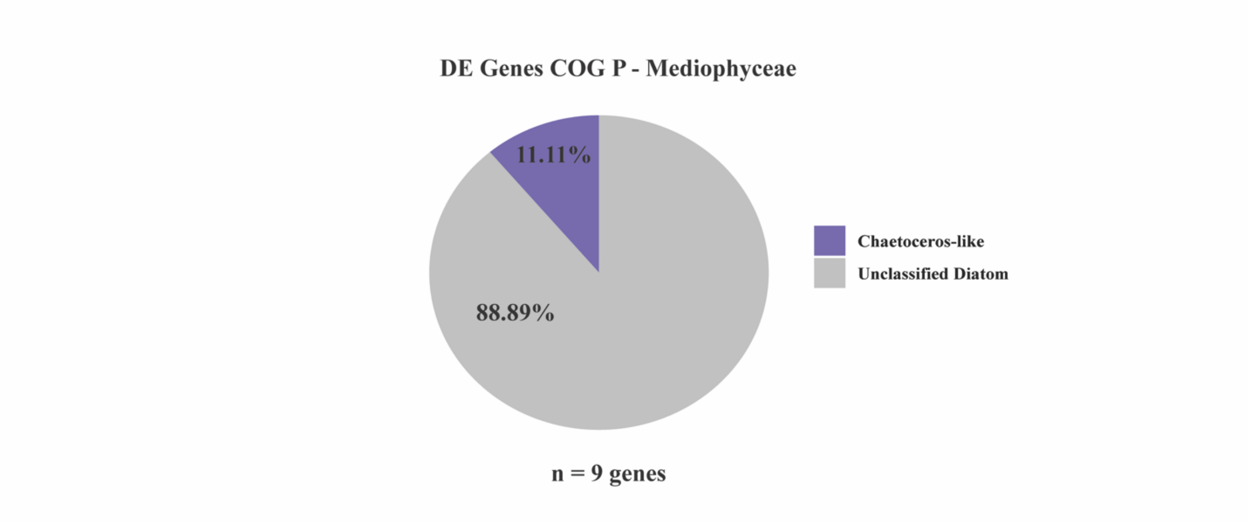


**Supplemental Figure 16:** Taxonomic distributions of Mediophyceae diatom genes that were differentially expressed by ice cover within COG Category P.

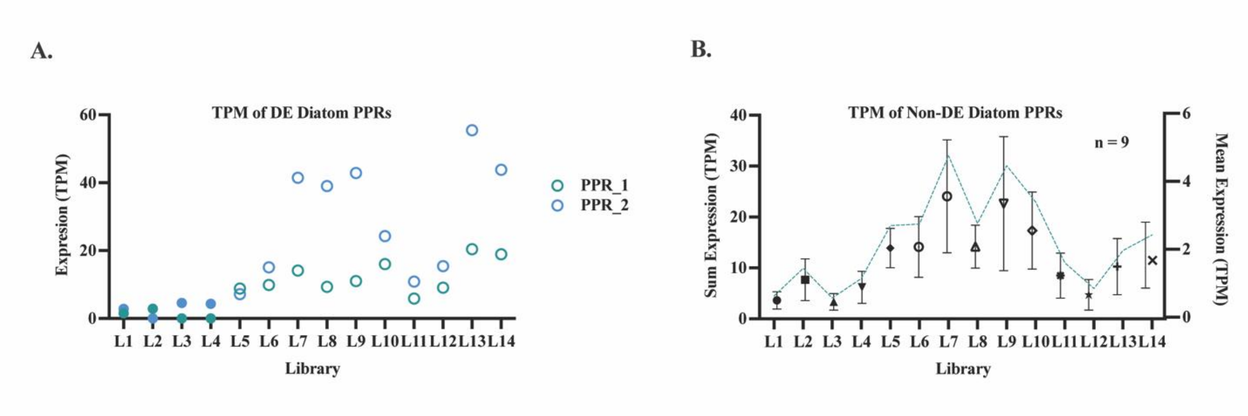


**Supplemental Figure 17:** Normalized expression (TPM) trends of diatom PPRs across winter libraries. (A) Normalized expression of the two DE diatom PPRs in ice cover (solid circles) and ice-free (open circles) winter libraries (B) Sum normalized expression of the 9 non-DE diatom PPRs in ice cover and ice free winter libraries indicated by the blue line (left y-axis). Mean expression + SEM of the 9 non-DE diatom PPRs in ice cover and ice free winter libraries indicated by the various shapes (right y-axis).

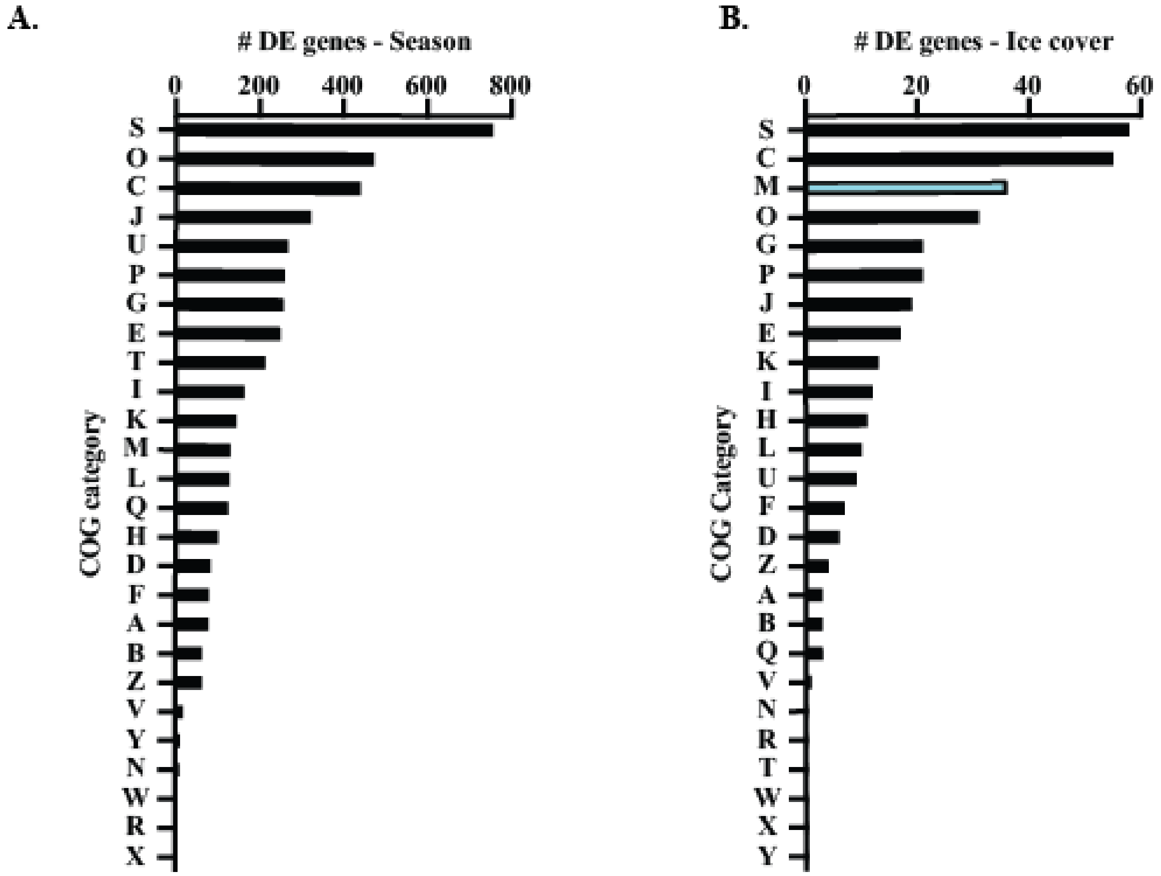


**Supplemental Figure 18:** Distribution of DE genes by COG category. **(A)** COG assignments for all DE genes (n = 4,854) in response to season (winter vs. spring). **(B)** COG assignments for all DE genes (n = 354) in response to ice cover, with COG category M indicated in blue.

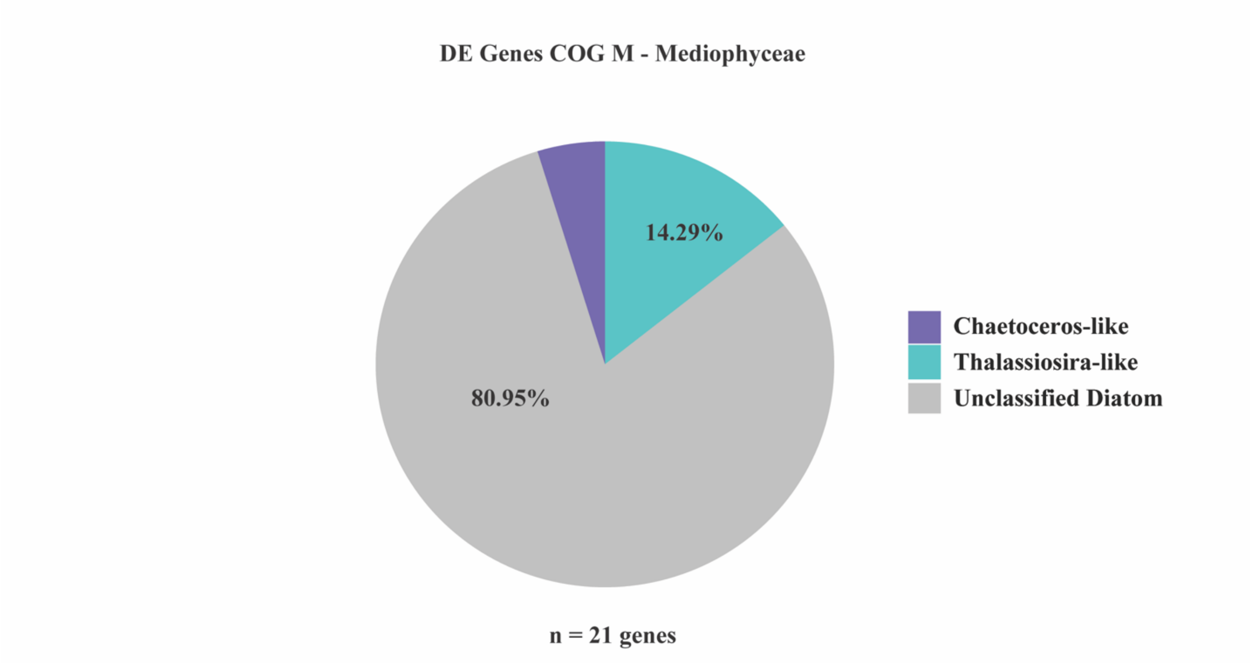


**Supplemental Figure 19:** Taxonomic distributions of Mediophyceae diatom genes that were differentially expressed by ice cover within COG Category M.

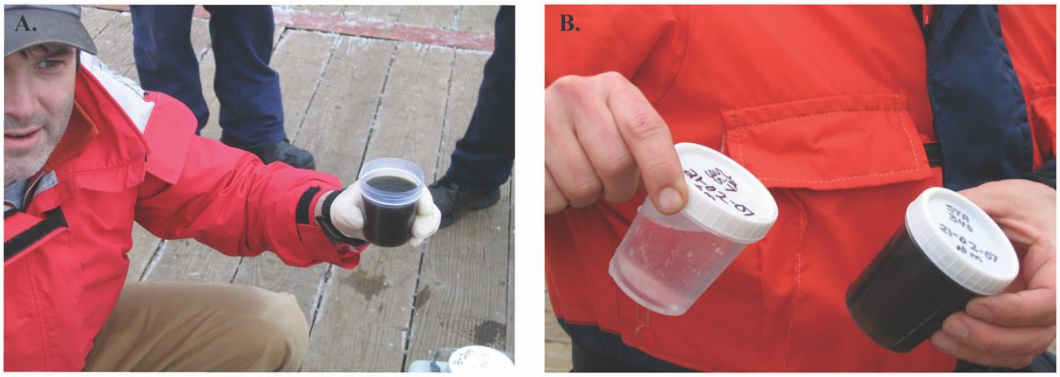


**Supplemental Figure 20:** Plankton net tows from a 2007 Lake Erie winter survey. (A) Concentrated diatom seston from a net-tow conducted in ice free waters at station 340 at 10 meters depth. (B) Dilute seston from a net-tow conducted in ice-covered waters at station 357 at 6 meters depth. Both samples were collected on February 2, 2007. Dr. R. Michael M. McKay is holding the samples, Dr. Steven W. Wilhelm took the photographs.
