## Supplemental Methods for "Declines in ice cover induce light limitation in freshwater diatoms"

For publication in conjunction with the following:

***Correspondence:**

**Supplemental Methods**

*Lake Erie winter-spring water column sampling*

To assess Chl *a* concentration, 50 mL volumes of sample water were collected on 0.22-$\mu$m nominal pore-size polycarbonate filters and stored at -20°C until Chl *a* extraction at *Bowling Green State University*. In addition, 200 mL volumes of sample water were collected on 20 $\mu$m nominal pore-size polycarbonate filters and stored at -20°C. Subsequently, samples were subject to a 24-hour extraction in 90% acetone at 4°C and quantified on a fluorometer (Turner Designs TD-700) equipped with a blue mercury bulb, a #10-050R excitation filter (340–500 nm) and a #10–115 (680 nm) emission filter [1]. Samples for RNA isolation were filtered through 0.22-$\mu$m nominal pore-size filters until the filter was saturated, flash frozen, and stored at -80 °C until extraction at *The University of Tennessee Knoxville*. For dissolved nutrients ($SiO_2$, $NO_3$, $NO_2$, $NH_3$, SRP, $SO_4$, Cl), 50 mL volumes of filtrate from the sterivex were collected and stored at -20°C until processing. Whole water samples for particulate nutrients (TN, TP) were also stored at -20°C until processing at *The Ohio State University Stone Laboratory.* There, samples were subsequently processed on a SEAL Analytical QuAAtro 5-Chaneel continuous segments flow auto-analyzer [2]. Fifty milliliter samples of whole water were preserved with Lugol's iodine and stored at room temperature until phytoplankton identification and enumeration at *Aquatic Taxonomy Specialists*, (Malinta OH). Briefly, samples were analyzed by counting phytoplankton in a measured aliquot using a modified inverted microscope for the Utermöhl method plus a small magnification modification of the stratified counting technique of Munawar and Munawar [3]. A measured aliquot of mixed sample was placed into an inverted microscope counting chamber and allowed to settle a minimum of 4 h per centimeter of overlying water depth. Larger and recognizable rare cells were counted at 400× along a minimum of one transect across the entire counting chamber. Smaller algae were counted at 1000× along a measured transect until a minimum of 300 cells were enumerated. Phytoplankton were counted as individual cells. All water column data was plotted in prism (v.9.3.1) by julian day to account for the interannual variability (2019-2020). Visualization of the 12 sample sites was performed in R using tmap [4] and open source shapefiles of Lake Erie shoreline (https://gis-michighan.opendata.arcgis.com/datasets) and bathymetry (https://www.ngdc.noaa.gov/mgg/fliers).

*Metatranscriptomic analysis*

The bioinformatic workflow used to process the libraries within this manuscript (Supplemental Table 1A) was initially compiled and published by Gilbert, LeCleir [5]. Trimming and filtering of raw reads to remove adaptors, contaminants, and sequence spike in-ins was performed by JGI *via* BBDuk (v.38.92) within the BBtools package [6]. Following, ribosomal RNA was removed using BBMap (default settings) (v.38.86) in line with the DOE JGI pipeline [7]. Filtered, trimmed reads from all 77 metatranscriptomic libraries were concatenated and assembled with MEGAHIT (v.1.2.9) [8], with the concatenated file serving as the input as performed previously [5]. Quality assessment of the coassembly was performed using QUAST (v.5.0.2) [9] with all statistics reported (Supplemental Tables). Trimmed, filtered reads from each library (n=20) were mapped to the coassembly *via* BBMap (default settings) (v.38.86) with mapping statistics reported (Supplemental Tables). Following, calling of the open reading frames (ORFs) within the coassembly was performed *via* MetaGeneMark (v.3.38) [10, 11] using the gene finding algorithm. Taxonomic annotation of the coassembly genes was performed using EUKulele [12] to perform a blast against the PhyloDB (v.1.076) database. Functional annotation of coassembly genes was conducted *via* eggNOG-mapper using orthology data and a specified e-value of $1e^{-10}$ (v.2.1.7) [13]. In this study, genes that were not assigned a "Preferred Name" by eggNOG were given one which we made based on the gene description provided by eggNOG-mapper. A list of the genes and their eggNOG preferred name (or the name we assigned in this study) is provided in the Supplemental Materials (Supplemental Tables). Trimmed, mapped reads were then tabulated according to ORF coordinates using featureCounts [14] within the subread (v.2.0.1) package. After, the tabulated read counts, EUKulele annotations, and eggNOG-mapper annotations were merged into a single file in R (v.4.0.0).

*Relative transcript abundance and taxonomy*

To investigate taxonomic trends inferred from proportional transcript abundance, coassembly genes were sorted according to the 7 classification levels of taxonomy assigned by EUKulele (domain-genus/species). To focus on diatoms specifically, analysis of eukaryotic reads mapped to major eukaryotic phytoplankton phyla (classification level 3 and 4), Bacillariophyta classes (classification level 6), and genus (classification level 7) were performed to determine relative proportions of the transcriptionally active community as performed previously (Supplemental Tables)[15]. Major eukaryotic phytoplankton taxa common to Lake Erie were selected based on prior long-term environmental surveys and manual sorting [16]. In the analysis of the relative transcript abundance, groups that were <5% of the total mapped reads were classified as "Other" as performed previously [15]. Due to the low resolution of taxonomic annotation at the genus level, "not annotated" reads were not included in the calculations at the genus level (Supplemental Figures 9-12). Rather, the unclassified diatom reads were included within the "Other" group to allow for the visualization of taxonomic trends of annotated taxa.

Relative abundances were graphically visualized in R using ggplot2 [17] and further modified in Adobe Illustrator (Adobe Inc., 2022).

In this study, we used the taxonomic annotation program EUKulele, which uses PhyloDB and the Marine Microeukaryote Metatranscriptomics Sequencing Project reference databases due to their unmatched depth and diversity. Hence, at the genus level all genera are reported as “genus-like” in this study to reflect that these are annotations made based largely on marine taxonomy which we have extrapolated to our freshwater system due to a lack of freshwater taxa represented within the databases. For example, *Aulacoseira*-like annotated genes within our dataset can be confidently suggested to be *Aulacosiera* based on freshwater cell abundances and taxonomy, yet we report it as *Aulacoseira*-like for consistency as it was annotated as the marine diatom *A. subartica*. Unfortunately, freshwater systems have not been as comprehensively sequenced/studied as marine systems. Indeed, TARA Oceans propelled the marine sequencing and taxonomic curation far beyond freshwater capacities [18]. Hence, this study illustrates the limitations within freshwater bioinformatic studies broadly.

### Results

#### *Taxonomically resolved winter diatom community transcriptional response to ice cover*

Bacillariophyta reads were dominated by the class Mediophyceae (polar centric diatoms), which remained the most abundant of the transcriptionally active winter diatom community regardless of ice cover (Figure 3B). Coscinodiscophyceae (centric) was the second most abundant, while pennate classes Bacillariophyceae and Fragilariophyceae remained at very low transcriptional abundances. Approximately 25% of the winter Mediophyceae community were classified as *Thalassiosir*a-like (Supplemental Figure 7), while 50% of the Coscinodiscophyceae winter community was annotated as *Aulacoseira*-like (Supplemental Figure 8). There were no clear trends within either centric class at the genus level. In contrast, pennate diatom classes Bacillariophyceae and Fragilariophyceae exhibited potential trends in genera transcript abundance as a function of ice cover, albeit minute (Supplemental Figures 9-10).

*Phylogenetic analysis of fasciclins*

The FAS1 domain appears to primarily originate from bacteria and is widely distributed throughout the bacterial domain. The FAS1 domain was found in 141 diatoms which span freshwater and marine distributions. Further, phylogenetic analysis demonstrates multiple instances of horizontal gene transfer (HGT), thus the FAS1 domain within diatoms appears to have been horizontally acquired from bacteria. Two of the putative HGT branches are quite long. This is likely due to lack of available genomes/sequencing for the respective diatom species. One long HGT branch corresponds to the diatom *Mayamaea pseudoterrestris,* the only member of genus *Mayamaea* with a genome sequenced (many of the individuals in this group are newly defined–*e.g.*, 2020, 2021, 2022). The second is from a transcriptomic dataset, and labeled as an unculturered *Nitzschia*. Four genomes are sequenced for this group, however given that it is an environmental sample it could likely be considered as "*Nitzschia"* because that is the closest relative/hit to current databases. Therefore, it could be a group not defined or not well defined to date. Generally, the branch lengths are long for the "diatom section", also likely related to incomplete taxonomy. Notably, the FAS1 domain is well represented in diatoms compared to other alga and cyanobacteria, yet most of the FAS1 diatom domains were annotated as "hypothetical, unnamed, or predicted protein" (Supplemental Tables). Hence, the FAS1 domain is not well-annotated or characterized within diatoms. Cumulatively, this data suggests the FAS1 domain distribution within diatoms is a result of HGT events with bacteria. Further, this data suggests the FAS1 domain is well-distributed across at least 141 diatom taxa as found in this study yet remains unannotated and undefined in most publicly available diatom sequencing data.
